# SQ3370, the first clinical click chemistry-activated cancer therapeutic, shows safety in humans and translatability across species

**DOI:** 10.1101/2023.03.28.534654

**Authors:** Sangeetha Srinivasan, Nathan A. Yee, Michael Zakharian, Maša Alečković, Amir Mahmoodi, Tri-Hung Nguyen, José M. Mejía Oneto

## Abstract

**Background:** SQ3370 is the first demonstration of the Click Activated Protodrugs Against Cancer (CAPAC™) platform that uses click chemistry to activate drugs directly at tumor sites, maximizing therapeutic exposure. SQ3370 consists of a tumor-localizing biopolymer (SQL70) and a chemically-attenuated doxorubicin (Dox) protodrug SQP33; the protodrug is activated upon clicking with the biopolymer at tumor sites. Here, we present data from preclinical studies and a Phase 1 dose-escalation clinical trial in adult patients with advanced solid tumors (NCT04106492) demonstrating SQ3370’s activation at tumor sites, safety, systemic pharmacokinetics (PK), and immunological activity.

**Methods:** Treatment cycles consisting of an intratumoral or subcutaneous injection of SQL70 biopolymer followed by 5 daily intravenous doses of SQP33 protodrug were evaluated in tumor-bearing mice, healthy dogs, and adult patients with solid tumors.

**Results:** SQL70 effectively activated SQP33 at tumor sites, resulting in high Dox concentrations that were well tolerated and unachievable by conventional treatment. SQ3370 was safely administered at 8.9x the veterinary Dox dose in dogs and 12x the conventional Dox dose in patients, with no dose-limiting toxicity reported to date. SQ3370’s safety, toxicology, and PK profiles were highly translatable across species. SQ3370 increased cytotoxic CD3^+^ and CD8^+^ T-cells in patient tumors indicating T-cell-dependent immune activation in the tumor microenvironment.

**Conclusions:** SQ3370, the initial demonstration of click chemistry in humans, enhances the safety of Dox at unprecedented doses and has the potential to increase therapeutic index. Consistent safety, toxicology, PK, and immune activation results observed with SQ3370 across species highlight the translatability of the click chemistry approach in drug development.

**Trial registration:** NCT04106492; 7 September 2019

## Introduction

For most therapies administered by systemic routes, only a small fraction of the drug ultimately reaches the target site in the body. The remaining dose is distributed elsewhere, free to engage in off-target interactions and induce adverse effects. Also, poor bioavailability of the drug at the target site can constrain how well it works [1–3]. Even the most promising therapies are often limited by their narrow therapeutic index (TI), which is a measure of the effective dose of a drug relative to its safety. Precisely targeting drugs to the site of interest can broaden their TI, thereby making them safer and more effective.

While preclinical pharmacological and toxicological data may be used to inform clinical outcomes of investigational drugs [4, 5], the TI estimated from these studies often does not translate to humans due to unmanageable toxicities, pharmacokinetic (PK) differences between species, or lack of efficacy [6]. Targeted approaches have been developed to overcome these limitations, but these have not been entirely successful because of inherent biological variability among *in vitro* studies, animal models, and human disease [7, 8]. Limited translatability across species may also be due to differences in biological characteristics of tumors required for drug activation, such as pH [9, 10], oxygen level [11, 12], protein expression [13], or enzymatic activity [14–17].

Click chemistry is a simple, powerful, Nobel prize-winning technology [18] with the potential to overcome the persistent barriers of unmanageable toxicities, PK differences between species, and biological variability in patients, while expanding the TI of drugs; leveraging this technology can accelerate drug development. Click chemistry involves chemical reactions in which two molecules react only with each other, ignoring any other molecules they encounter. Click chemistry reactions that occur in a biological environment are termed bioorthogonal chemistry [19]. As the molecules only react with each other during click chemistry activation, there is no interference from or to native biological processes.

Shasqi Inc. (San Francisco, CA, USA) is pioneering click chemistry in humans with a focus on cancer therapeutics. Cancer is a major challenge worldwide, with 2020 estimates of 20 million new cancer cases and 10 million cancer deaths [20]. The toxicities associated with cancer treatments are significant and can often result in poor quality of life and even death [21]. While currently available targeted therapies often have a more favorable toxicity profile, many still have undesirable off-target effects and are limited by the biology of the tumor [22]. Some cancer treatments also suppress the immune system, lowering the body’s ability to fight cancer [23], while others overstimulate the immune system, leading to autoimmune disease or cytokine release syndrome [24]. These off-target toxicities limit the anti-neoplastic activity of cancer therapeutics in the clinic, narrowing the TI. The objective of Shasqi’s Click Activated Protodrugs Against Cancer (CAPAC™) platform is to provide an effective mechanism, based on click chemistry, to activate cancer therapies directly in the tumor and its microenvironment.

The CAPAC platform consists of two components: a tumor-localizing agent and a chemically attenuated cancer drug (protodrug). When the two components come together in a biological environment, they “click” through an irreversible covalent reaction of the moiety on the localizing agent and a moiety on the protodrug, leading to the release of the active payload into the tumor and its microenvironment.

Doxorubicin (Dox) is an anthracycline chemotherapeutic agent that has been widely used for over 40 years to treat solid tumors of diverse types [25]. Toxicities such as myelosuppression limit the Dox dose per cycle and late effects such as cardiomyopathy and leukemogenesis limit the Dox total dose that a patient can receive over a lifetime [26, 27]. Targeting approaches over the last 30 years have largely failed to overcome these known Dox toxicities as they relied on tumor biology such as enhanced permeability and retention, enzyme expression, and different physiological characteristics [9, 25, 28], which are subject to variations across tumor types and patients. Thus, Dox is a good candidate for CAPAC, and localization of high concentrations of Dox to tumors has the potential to fulfill an unmet clinical need.

SQ3370 is the first click-activated investigational therapy developed, and consists of: 1) a tumor-localized SQL70 biopolymer, which is a derivative of sodium hyaluronate chemically modified with tetrazine, and 2) the SQP33 protodrug, which consists of a Dox payload whose cytotoxic activity is chemically attenuated by modification with *trans*-cyclooctene [28]. When the SQP33 protodrug meets the SQL70 biopolymer at the tumor site, a rapid click reaction occurs between the tetrazine moiety of SQL70 and the *trans*-cyclooctene moiety of SQP33 (the click chemistry groups with complementary reactivity), selectively releasing Dox at the tumor and its microenvironment (Fig. 1).

**Fig. 1.**
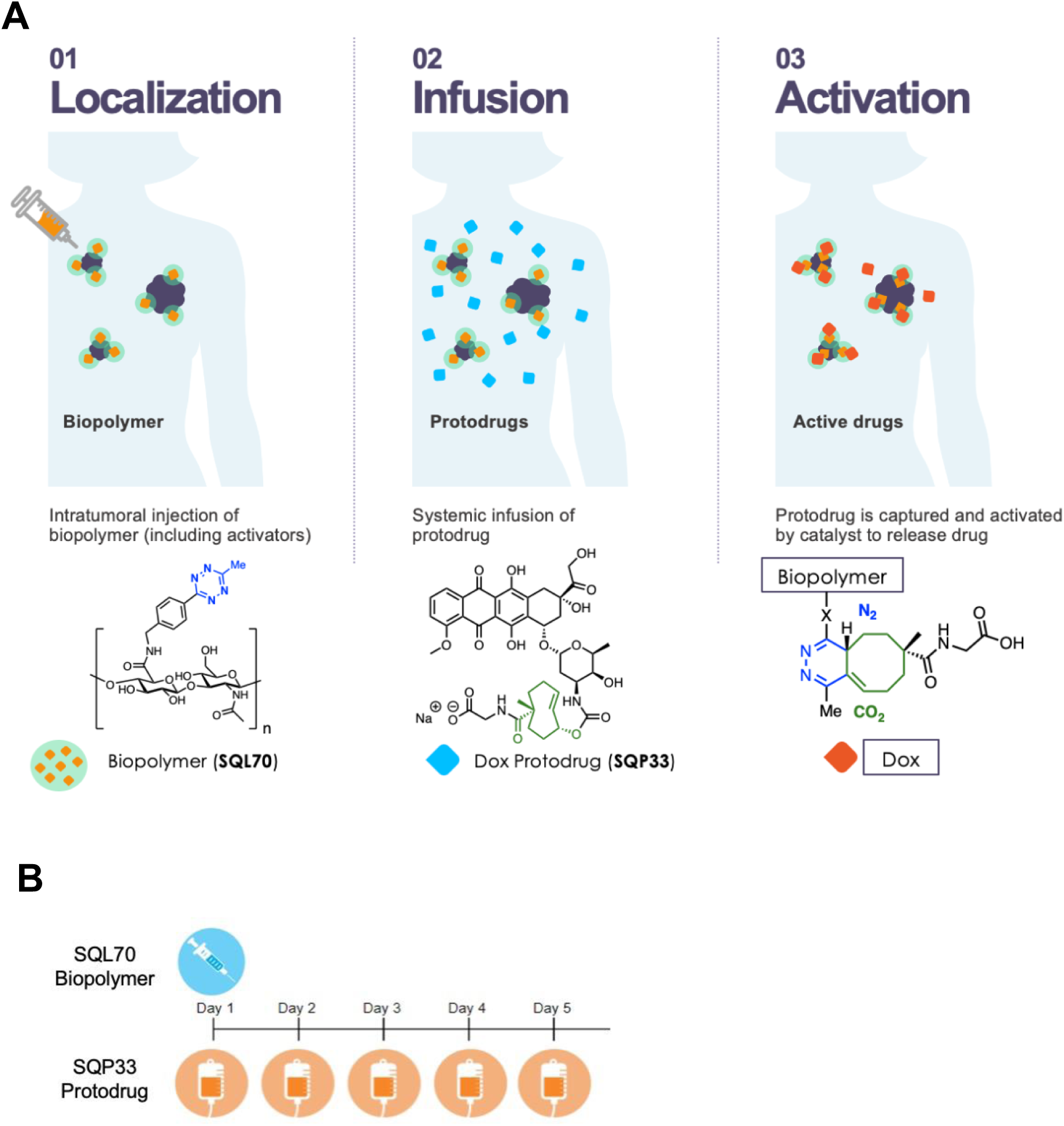
Overview of SQ3370 local tumor targeting with CAPAC platform. (**A**) 1) SQL70 biopolymer is locally injected at the tumor site. 2) SQP33 protodrug is infused systemically. 3) SQP33 protodrug is activated by SQL70 biopolymer through a rapid covalent reaction between tetrazine and *trans*-cyclooctene moieties, followed by chemical rearrangement to release active Dox. (**B**) The standard dosing regimen of SQ3370 includes 5 daily systemic IV infusions of SQP33 protodrug beginning 1 to 4 hours after SQL70 biopolymer injection on day 1. In SQ3370-001, a Phase 1/2a clinical trial in patients with advanced or metastatic solid tumors, SQ3370 is administered in 21-day cycles, with SQL70 biopolymer injected at the local tumor site on day 1, followed by protodrug doses on days 1 to 5. CAPAC = Click Activated Protodrugs Against Cancer; Dox = doxorubicin; SQ3370 = SQP33 protodrug + SQL70 biopolymer; IV = intravenous.

We have previously reported on the safety, tolerability, PK, and efficacy of SQ3370 in rodents [28, 29]. SQ3370 was well tolerated at 19.1-times (mice) and 10.8-times (rats) the maximum tolerated dose (MTD) of conventional Dox in the respective species (Dox MTD: 20 mg/kg in mice and 2.5 mg/kg in rats). SQP33 protodrug was stable *in vivo* showing negligible spontaneous activation in the absence of the SQL70 biopolymer [29]. In healthy rats, SQ3370 significantly lowered Dox exposure to the heart, liver, and kidney compared with conventional Dox treatment [29]. SQL70 biopolymer remained at the subcutaneous (SC) injection site for an extended duration and could effectively activate multiple IV doses of SQP33. Further, in a dual tumor model of MC38 tumors, where only one lesion was injected with the biopolymer, SQ3370 enhanced anti-tumor responses against the injected and non-injected lesions, increasing overall survival by 63% over conventional Dox treatment [28]. The response in the non-injected lesions is due to an increase in immune activation in the form of CD4^+^ and CD8^+^ T-cells that was observed in the tumors of SQ3370-treated mice. These encouraging results paved the way for the clinical development of SQ3370, which is currently being evaluated in a Phase 1/2a clinical trial in patients with advanced or metastatic solid tumors (SQ3370-001; NCT04106492).

In this article, we report on the ability of the CAPAC platform to target tumors with high concentrations of Dox using the click chemistry-activated SQ3370 as a model. We present data demonstrating SQ3370’s activation at tumor sites, safety, systemic PK, and immunological activity from preclinical studies and a Phase 1 dose-escalation clinical trial in adult patients with advanced solid tumors. We highlight the translatability of click chemistry across species and utility in the discovery of new biological insights (immune activation) for the development of potential new cancer therapeutics.

## Materials and Methods

### Preclinical studies

All the procedures related to animal handling, care, and treatment were preformed according to the guidelines approved by the Institutional Animal Care and Use Committee (IACUC) of each institution, following the guidance of the Association for Assessment and Accreditation of Laboratory Animal Care (AAALAC).

#### Efficacy of SQ3370 in a single tumor MC38 mouse model

The colorectal carcinoma MC38 murine model studies were conducted at BioDuro (at Explora BioLabs’ vivarium in San Diego, CA, USA). The MC38 cell line was obtained from Kerafast (Boston, MA, USA; cell line originated from the National Cancer Institute/National Institutes of Health) and was used to establish the MC38 mouse model [30]. The cells were expanded in complete culture medium RPMI-1640 containing 20% FBS, 2 mM glutamine, and 1% Pen-Strep in a 37 °C incubator in an atmosphere of 5% CO_2_. The cells were routinely passaged twice a week. Once the MC38 cells reached the exponential growth phase, the cells were harvested, washed, and counted for cell inoculation in mice. Forty female C57BL/6 mice were inoculated with 5 x 10^5^ MC38 tumor cells via SC inoculation in the right flank and the tumors left to develop. When the tumors reached approximately 75 mm^3^, 20 mice received 100 µL of SQL70 via a peritumoral (*n* = 10) or SC injection in a distal site in the left flank (*n* = 10) on day 1, followed by five daily IV injections of SQP33 protodrug at a previously established MTD schedule (383 mg/kg/cycle Dox Eq, beginning day 1 within one hour of receiving SQL70). Dox (single dose on day 1 at 20 mg/kg) and saline control groups did not receive SQL70 biopolymer injections. Tumor volumes were measured twice weekly until tumors reached at least 2,000 mm^3^. To assess Dox exposure and necrosis at the tumor site, in another MC38 tumor study conducted at Cephrim Biosciences (Woburn, MA, USA), mice were dosed with Dox (IV, QDx2, 8.1 mg/kg/dose) or SQ3370 [SQL70 intratumorally with SQP33 (IV, QDx2 or QDx5, 78.6 mg/kg/dose Dox Eq)], SQP33 alone (IV, QDx2, 78.6 mg/kg/dose Dox Eq) or saline. Tumors were isolated one hour after the last IV dose and the tumor tissue was processed into thin sliced sections, which were subjected to either hematoxylin and eosin (H&E) staining or preparation for MALDI imaging assays. Additional details can be found in the supplementary methods.

#### GLP toxicology and TK evaluation in dogs

A GLP study to determine the toxicology and TK effects of SQ3370 (including SQP33 protodrug, Dox, and SQL70 biopolymer) treatment in dogs was conducted at Covance Laboratories (Madison, WI, USA). Dogs were selected for toxicological testing as they are more sensitive to the cardiotoxic effects of Dox than other species, including humans [31]. Dose selection was informed by the results of a non-GLP study conducted in non-naïve dogs that showed the MTD in dogs is 14 mg/kg/cycle Dox Eq (data not shown).

Seventy-two male and female purebred Beagle, healthy, treatment-naïve dogs were assigned to 8 groups, using a procedure to balance mean weight and sex across groups. Three experimental groups received SQ3370, made up of a single SC biopolymer injection of 10 mL on day 1 and daily IV infusions of the SQP33 protodrug on days 1-5, according to their assigned group dose level (1.8, 3.6, or 8.9 mg/kg/cycle Dox Eq). The effects of SQP33 protodrug alone (without the drug-activating biopolymer) were assessed at the highest dose level (8.9 mg/kg/cycle Dox Eq) in one group with a vehicle control SC injection of 10 mL. The effects of SQL70 biopolymer alone (without any functionally active drug payload) were evaluated at 3 dose volumes: 5 mL, 10 mL, and 15 mL (given as 7.5 mL x 2 SC injections). One group received a vehicle control for both the SQP33 protodrug and the SQL70 biopolymer.

Assessment of toxicity was based on clinical observations, body weights and weight fluctuations, food consumption, ophthalmic observations, ECG measurements, clinical and anatomic pathology, and mortality throughout the 5-day dosing period and a 23-day recovery phase. The anatomic pathology evaluations consisted of macroscopic observations, the weighing of selected organs, and the histopathological examination of a full list of tissues from all animals.

Blood was drawn 5 min, 30 min, 1 h, 3 h, 8 h, and 24 h following SQP33 infusion on days 1 and 5, and liquid chromatography-tandem mass spectrometry (LC-MS/MS) was used to measure plasma SQP33 protodrug and Dox concentrations. Plasma bioanalysis was conducted at Covance Laboratories (Salt Lake City, UT, USA).

### Clinical evaluation

#### Study design and patients

This single arm, open-label, first-in-human Phase 1/2a study ([NCT041064926](https://clinicaltrials.gov/ct2/show/NCT04106492)) was conducted at multiple sites across the United States and Australia. The study was approved by the Institutional Review Boards at each site and conducted according to the Declaration of Helsinki. Patients provided written informed consent before any study procedures were performed.

Male and female patients of at least 18 years of age with locally advanced or metastatic solid tumors with a lesion amenable to repeated biopolymer injection and for which anthracycline-containing treatment was indicated were eligible to enroll. Patients were either relapsed or refractory following standard of care therapy, or were ineligible for standard of care, and must have had ≤ 300 mg/m^2^ prior Dox exposure.

Eligible patients were enrolled into a series of SQP33 protodrug dose escalation cohorts consisting of a Phase 1 accelerated single cohort design, followed by transition to a Phase 2 Rolling 6 cohort design. The study treatment was delivered in 3-week (21-day) cycles with SQL70 biopolymer administered via peritumoral injection on day 1, followed by 5 daily infusions of SQP33 protodrug on days 1–5. SQL70 biopolymer was administered in a 10 mL or 20 mL injection volume, in up to two lesions and/or defined clusters of lesions. Patients could continue on treatment until they had no injectable lesions, disease progression, unacceptable toxicity, or other treatment discontinuation criteria were met. The enrollment for the Phase 1 dose escalation trial has ended. However, as some patients in the 15x dose cohort continue to receive treatment, data from this cohort is not presented here. Further, data from patients who received 20 mL biopolymer injections is not reported here.

#### Pharmacokinetics

Blood was collected from subjects on each day of dosing to characterize the plasma PK of SQP33 protodrug and Dox. Plasma samples were processed using solid-phase extraction followed by analysis with high performance liquid chromatography-tandem mass spectrometry (LC-MS/MS). Clinical plasma bioanalysis was conducted at Agilex Biolabs (Thebarton, SA, Australia).

#### Sequential immunohistochemistry, image acquisition, and processing

Human tissues used for immunohistochemical analysis were retrieved and fixed with 10% buffered formalin, dehydrated in ethanol, and embedded with paraffin. Sequential immunohistochemistry (IHC) was performed on 5 μm formalin-fixed paraffin embedded (FFPE) sections using an adapted protocol based on methodology we previously described [32]. Briefly, slides were deparaffinized and stained with hematoxylin (S3301, Dako, Santa Clara, CA, USA), followed by whole-slide scanning at 20X magnification on an Aperio AT2 (Leica Biosystems, Wetzlar, Germany). Sectioned tissues then underwent 20 min heat-mediated antigen retrieval in pH 6.0 Citra solution (BioGenex, Fremont, CA, USA), followed by 10 minutes blocking in Dako Dual Endogenous Enzyme Block (S2003, Dako, Santa Clara, CA, USA) (human), then 10 min protein blocking with 5% normal goat serum and 2.5% BSA in TBST. For staining, primary antibody incubations were carried out for 1 hour at 4 °C using CC3 (ASP175, 1:400, Cell Signaling); and 30 min at room temperature using CD45 (H130, 1:100, E Bio), CD3 (SP7, 1:150, Thermo Sci), PANCK (AE1/AE3, 1:2,000, Cell Signaling), CD8 (SP16, 1:100, Invitrogen/Thermo), or GRZB (Polyclonal, 1:200, Abcam). After washing off primary antibody in TBST, either anti-rat, anti-mouse, or anti-rabbit Histofine Simple Stain MAX PO horseradish peroxidase (HRP)-conjugated polymer (Nichirei Biosciences, Tokyo, Japan) was applied for 30 min at room temperature, followed by AEC chromogen (Vector Laboratories, Burlingame, CA, USA). Slides were digitally scanned following each chromogen development, and the staining process was repeated for all subsequent staining cycles.

Scanned images were registered in MATLAB version R2020b using the SURF algorithm in the Computer Vision Toolbox (The MathWorks, Inc., Natick, MA, USA). Image processing and cell quantification were performed using FIJI (FIJI Is Just ImageJ) [33], CellProfiler Version 3.1.9 [34], and FCS Express 7 Image Cytometry RUO (De Novo Software, Glendale, CA, USA). AEC signal was extracted for quantification and visualization in FIJI using a custom macro for color deconvolution. Briefly, the FIJI plugin Color_Deconvolution [H AEC] was used to separate hematoxylin, followed by postprocessing steps for signal cleaning and background elimination. AEC signal was extracted in FIJI with the NIH plugin RGB_to_CMYK. Nuclear stain images were segmented in FIJI using the StarDist plugin [35]. Color deconvoluted images and segmented nuclei were processed in CellProfiler to quantify single cell mean intensity signal measurements for every stained marker, and cell populations were identified and quantified by image cytometry in FCS Express based on hierarchical expression of known markers. For visualization, stained images were color-deconvoluted, registered, and overlaid in HALO (Indica Labs, Albuquerque, NM, USA).

### Statistical analysis

Statistical analyses for nonclinical studies were performed using GraphPad Prism 9.0 (GraphPad Software, Boston, MA, USA). For PK samples in dogs, *p*-values were calculated using Welch’s *t*-test for comparisons of two groups and one-way analysis of variance (ANOVA) with Tukey’s post-test for comparison of three groups. For tumor volumes in MC38-bearing mice, *p*-values were calculated using the mixed effects model. Survival analysis in mice was performed using the Mantel-Cox logrank test and logrank Hazard ratios. *P*-values for analysis of mIHC staining in patient tumor biopsies from the clinical trial were performed using the Mann-Whitney *U* test. All other clinical data listings, figures and summary tables for the baseline characteristics, study drug administration, safety, and efficacy data were assessed with SAS software (9.4 or later). All listings were sorted by type of dosing schedule, dose level cohort, subject number, and assessment date/time. Continuous variables were summarized using descriptive statistics including number of non-missing observations, mean, standard deviation, median, and minimum and maximum values. Categorical variables were summarized with frequency counts and percentages.

## Results

### Preclinical evaluation

#### SQ3370 induces anti-tumor responses by localizing Dox to tumors in the MC38 mouse tumor model

Significantly greater anti-tumor responses were observed in MC38 tumor-bearing mice receiving SQL70 biopolymer directly at the tumor site than in mice in which SQL70 biopolymer was injected at a distal site (*p* = 0.0020 on day 20) or in mice that received conventional Dox (*p* = 0.0371 on day 17) (Supplementary Figure S1A). Median overall survival was also significantly improved in mice receiving biopolymer at the tumor site (31 days) compared with that in mice in which the biopolymer was injected at a distal site (24 days, *p* = 0.0015) or mice that received conventional Dox (25.5 days, *p* = 0.0031) (Supplementary Figure S1B-C), indicating that localization of the biopolymer at the tumor site plays an important role in disease control. Of note, anti-tumor response and overall survival with distal SQ3370 treatment were similar to those with Dox control treatment.

To further elucidate the intratumoral distribution and effects of Dox, matrix-assisted laser desorption/ionization-mass spectrometry imaging (MALDI-MSI) was used for tissue-based analysis of Dox in target lesions of MC38 tumor-bearing mice. Mouse tumor sections sampled after SQ3370 treatment showed high levels of Dox exposure (up to 101 µg/g of tissue) in the tumor and stroma on days 2 and 5 (Fig. 2). Histological assessment of these sections showed that these high levels of Dox correlated with notable areas of early and advanced necrosis. In contrast, tumor sections after conventional Dox treatment showed no detectable levels (limit of detection = 9.1 µg/g of tissue) of intratumoral Dox or advanced necrosis. Tumor sections from mice dosed with vehicle or SQP33 alone also did not result in any detectable level of Dox in the tumor (Supplementary Figure S2). Particularly, this study highlighted that SQ3370 can effectively activate the therapeutic Dox payload within the tumor at levels that are unachievable by conventional treatment approaches.

**Fig. 2.**
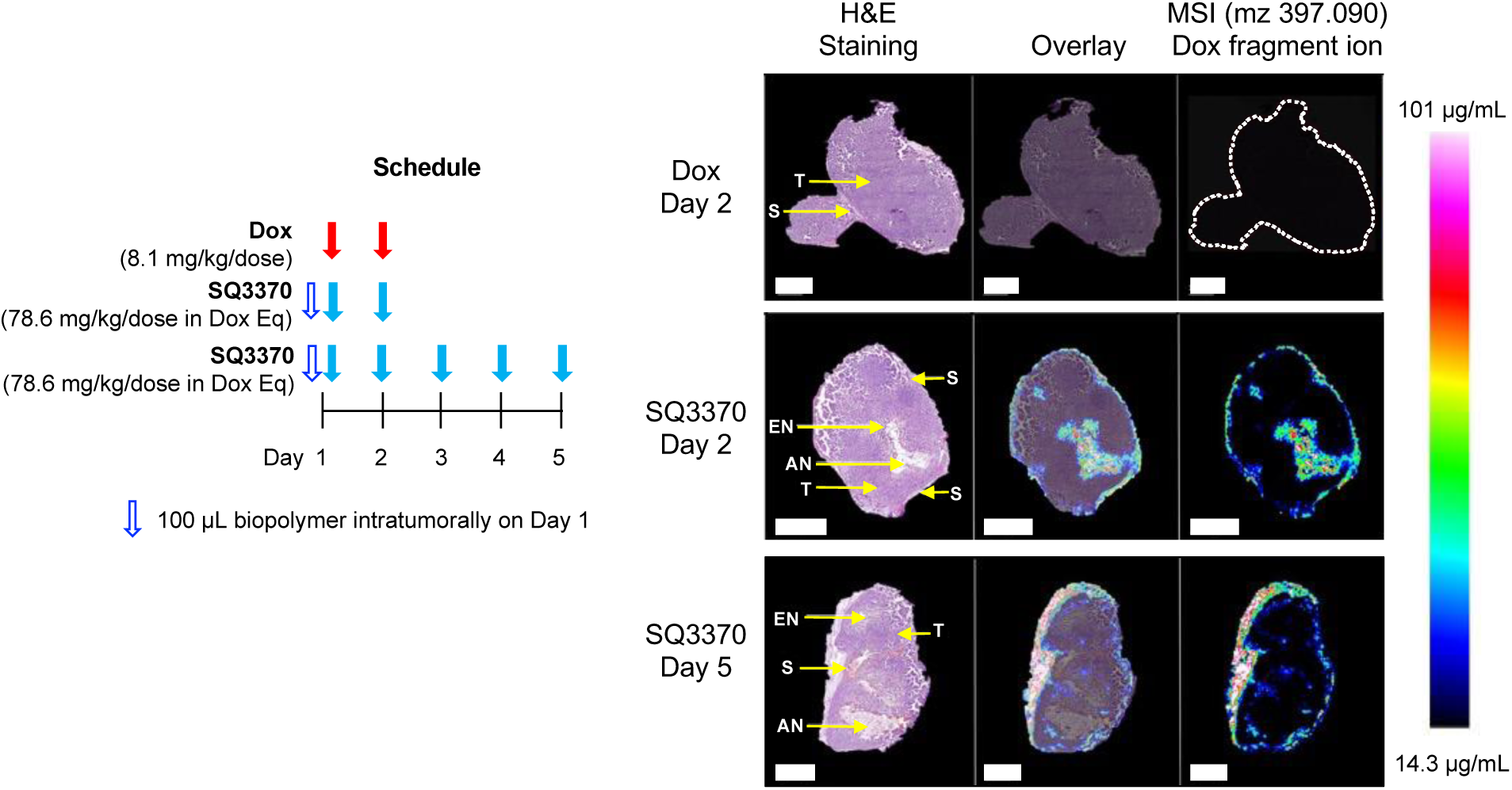
Tumor Dox activation and distribution after SQ3370 treatment in a mouse tumor model. Tumor tissues were collected from mice engrafted with MC38 tumor 1 hour after dosing with Dox (IV, QDx2, 8.1 mg/kg/dose) or SQ3370 [SQL70 intratumorally, SQP33 (IV, QDx2 or QDx5, 78.6 mg/kg/dose Dox Eq)]. Serial sections were generated on cryostat for MALDI-MSI and H&E staining. MALDI-MSI (right column) of SQ3370-treated tumors detected high Dox levels on days 2 and 5 that correlated with necrosis (H&E staining, left column), while conventional Dox-treated tumors did not. Scale bars: 2 mm. Limit of detection = 9.1 µg/mg of tissue. AN = advanced-stage necrosis; Dox Eq = Dox molar equivalents; EN = early-stage necrosis; H&E = hematoxylin and eosin; IV = intravenous; MALDI-MSI = matrix assisted laser desorption/ionization mass spectrometry imaging; MSI = mass spectrometry imaging; N = necrosis; S = stroma; T = tumor.

#### SQ3370 is safe and well tolerated in a Good Laboratory Practice (GLP) dog toxicology study

In naïve non-tumor bearing dogs, SQL70 biopolymer was given as a SC injection and was followed by 5 daily doses of SQP33 protodrug and a 23-day recovery period. The highest non-severely toxic dose (HNSTD) of SQ3370 was 8.9 mg/kg/cycle Dox molar equivalents (Dox Eq), which is 8.9x the standard veterinary dose of Dox in dogs (1 mg/kg/cycle) [36]. The dose-limiting toxicities (DLTs) observed at the HNSTD were consistent with those previously reported for conventional Dox [37], including moderate bone marrow hypocellularity and decreased hematopoiesis. SQP33 and SQL70 were well tolerated alone or together as SQ3370, with no treatment-related effects on ophthalmic or macroscopic observations or alterations in body weight, food consumption, or organ weight (Supplementary Results). Notably, there was no SQ3370-related cardiotoxicity in dogs as reported by electrocardiogram measurements. Dox is a recognized vesicant and can cause extensive tissue damage and blistering if it escapes from the vein [37]. However, SQ3370 did not induce a vesicant effect, consistent with activation of the Dox payload at the biopolymer injection site rather than an off-target infusion site. SQP33 protodrug given alone at 8.9 mg/kg/cycle Dox Eq, without biopolymer, resulted in no observable adverse effects. SQL70 biopolymer given alone, without the protodrug, resulted in an injection site reaction consistent with an inflammatory response to foreign material, consistent with lower molecular weight hyaluronic acid-based materials [38],without notable tissue irritation. Interestingly, these inflammatory effects were not seen for SQ3370. Overall, all observed reactions were reversible during the recovery period.

#### SQ3370 leads to efficient and persistent conversion into active Dox in GLP dog toxicokinetic (TK) assessments

Maximum plasma concentration (C_max_) resulting from SQ3370 treatment at the HNSTD and SQP33 alone (without SQL70 biopolymer) are shown in Fig. 3. Capture of the protodrug by the biopolymer and conversion to Dox was evident on both days 1 and 5. On day 1, the C_max_ of SQP33 was significantly reduced in the presence of SQL70 biopolymer (*p* < 0.0001, Fig. 3A), with an associated significant increase in plasma Dox (*p* < 0.0001, Fig. 3B), indicating efficient capture and activation. This is also reflected in the plasma concentration-time curves (Supplementary Figure S3A-B); SQP33 was undetectable in plasma after the 5-minute timepoint due to rapid capture and activation by the biopolymer. In the absence of biopolymer, SQP33 had a plasma half-life of slightly over 0.6 hours.

**Fig. 3.**
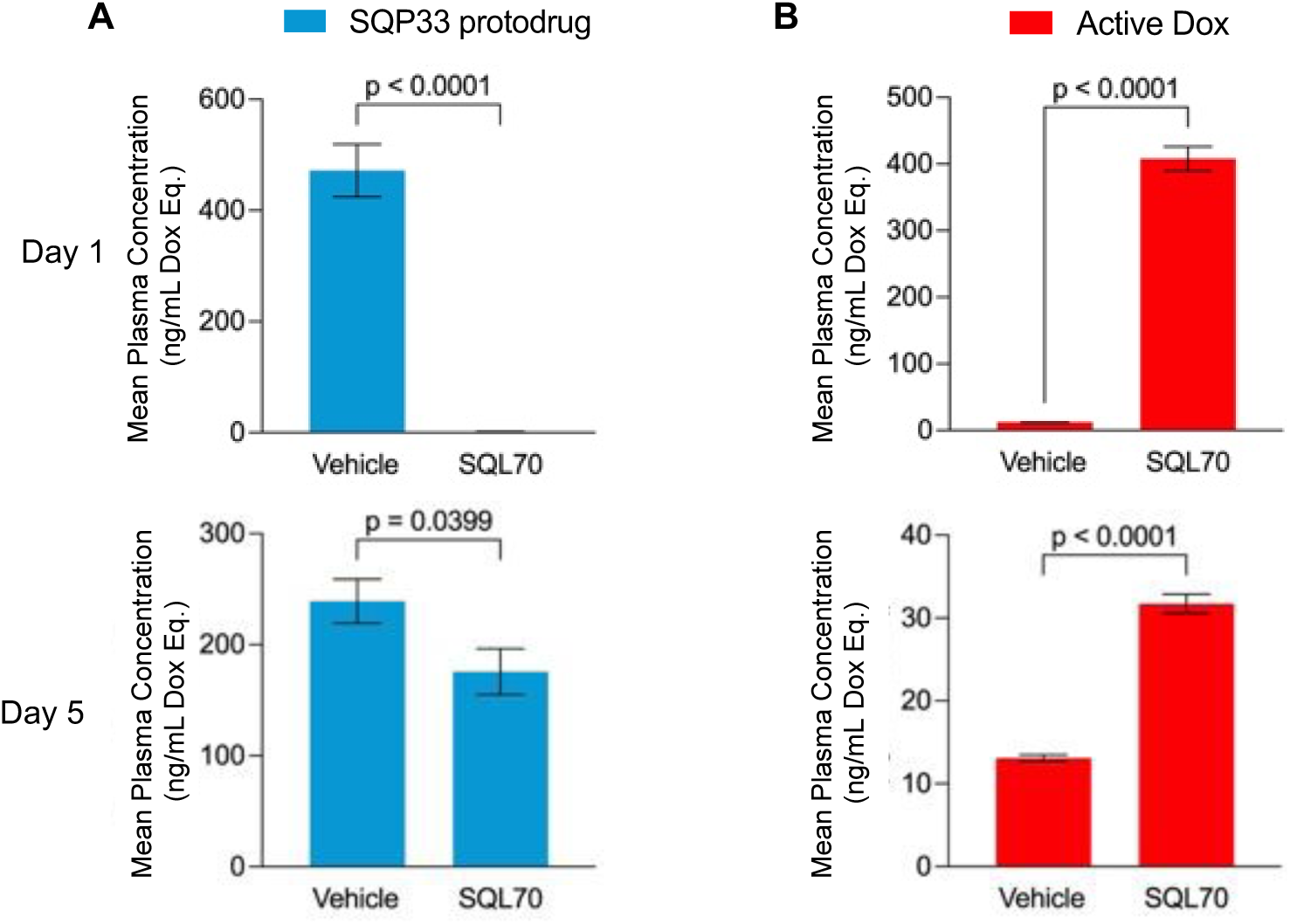
Maximum concentrations (C_max_) of SQP33 protodrug and active Dox in dogs on days 1 and 5. (**A**), (**B**) Male and female purebred Beagle dogs were treated with 8.9 mg/kg/cycle SQP33 (Dox Eq) over a 5-day dosing period (*n* = 20). On day 1, one group received 10 mL of SQL70 while the second group received 10 mL vehicle SC. Approximately 1 hour following SC injection, animals were administered SQP33 protodrug for approximately 30 minutes via IV infusion at a volume of 5 mL/kg/day. Plasma concentrations of SQP33 protodrug and active Dox were measured using LC-MS at 5 min, 30 min, 1, 3, 8 and 24 hours after SQP33 protodrug infusion on days 1 and 5. Maximum day 1 and day 5 active SQP33 **(A)** and active Dox concentrations **(B)** were compared relative to the respective vehicle group. Shown are mean ± SEM. Statistical significance was determined by two-tailed Welch’s t-test. Conc. = concentration; Dox = doxorubicin; Dox Eq = doxorubicin molar equivalents; IV = intravenously; LC-MS = liquid chromatography-mass spectrometry; SC = subcutaneous; SEM = standard error of mean.

On day 5, the SQP33 C_max_ in the presence of the biopolymer was reduced by 26.5% relative to the C_max_ in the presence of SQP33 alone (*p* = 0.0399), with a corresponding reduction in overall plasma exposure as seen on the time curve (Fig. 3A). SQP33 persisted longer on day 5 than on day 1, being detectable up to the 1-hour timepoint (Supplementary Figure S3A-B). These observations confirm that the biopolymer retained activating capacity by day 5, although it decreased over the course of the 5-day regimen. This finding was correlated with an increase in the average plasma Dox C_max_ on day 5 (*P* < 0.0001, Fig. 3B).

The plasma concentrations of SQP33 and Dox were dependent on the presence of the biopolymer. In animals treated with SQ3370, the average plasma Dox C_max_ values were approximately 33-fold and 2.4-fold higher than in those in animals treated with SQP33 alone on day 1 and day 5, respectively (Fig. 3B). Conversely, the average SQP33 C_max_ following SQ3370 treatment was 126-fold and 1.4-fold lower than those treated with SQP33 alone on day 1 and day 5, respectively (Fig. 3A). Together, these data also show the stability of SQP33 *in vivo* as it resulted in minimal conversion to Dox in the absence of the biopolymer. Furthermore, the Dox area under the curve (AUC) and C_max_ were found to be dose proportional, highlighting that Dox exposure increases with higher doses of SQ3370 (Supplementary Figure S3C-D). Overall, the observations in the dog study were consistent with those previously reported in rodents [29].

### Phase 1 clinical evaluation in patients with advanced solid tumors

#### Patient cohorts and baseline characteristics

Based on the encouraging preclinical data, the investigation of SQ3370 advanced to a first-in-human Phase 1/2a clinical trial. SQ3370-001 (NCT04106492) is a single-arm, open-label study conducted at multiple clinical sites in the United States and Australia. The design includes a Phase 1 dose escalation with the primary endpoint to determine the recommended Phase 2 dose (RP2D) of SQ3370, based on an evaluation of safety and tolerability, followed by a Phase 2a evaluating safety, tolerability, and efficacy in Dox-sensitive tumors. Patients received SQL70 biopolymer injected intratumorally into a single lesion on day 1 and SQP33 IV infusion once daily on days 1-5 (Figure 1B) of the 21-day treatment cycle.

The Phase 1 study enrolled 39 patients at nine escalating SQP33 dose levels, from 0.38x to 15x the conventional Dox dose of 75 mg/m^2^, per cycle. The study did not reach an MTD and dose escalation was halted at the 15x dose level. With no reported protocol-defined DLTs, the 12x dose was chosen as the RP2D. In this manuscript, we present data from the first 23 patients in the escalation cohorts up to the 12x dose level. (Fig. 4).

**Fig. 4.**
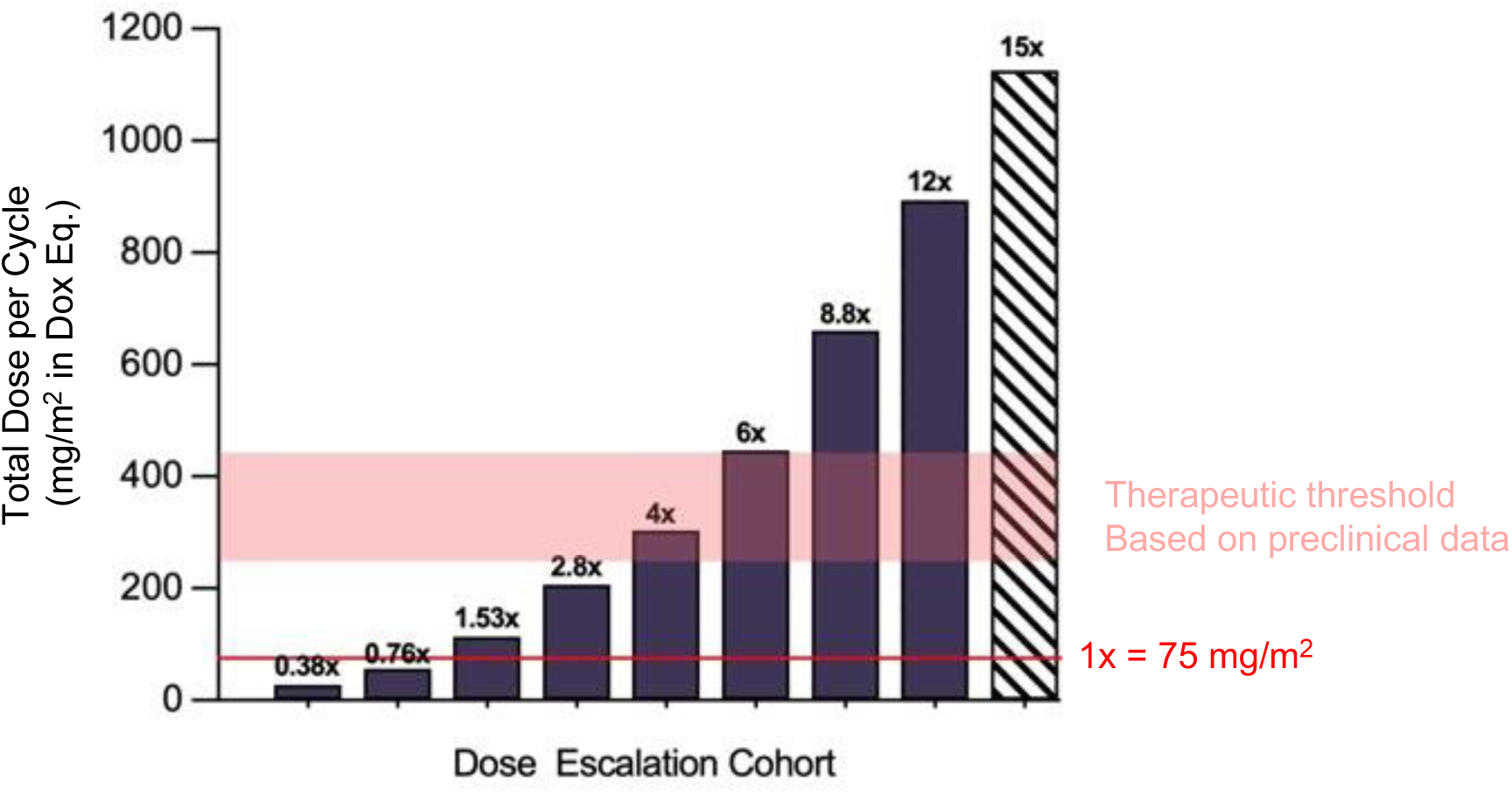
SQ3370-001 patient dose escalation. Human patients were dosed up to 15x the conventional dose of Dox for Phase 1 clinical trial dose escalation. At 15x the conventional dose of Dox (MAD), the dose escalation was halted, as the MTD had not been reached. The RP2D was established at 12x. Data is reported in this article up to the 12x dose level cohort. Conventional Dox dose (1x) = 75 mg/m^2^. Dox = doxorubicin; Dox Eq = doxorubicin molar equivalents; MTD = maximum tolerated dose; MAD = maximum administered dose; RP2D = recommended Phase 2 dose.

Patients were mostly white (87%), male (48%) with a median age (range) of 59 years (26–92) and were predominantly diagnosed with sarcoma (65%) (Table 1). Most had received prior systemic anti-cancer therapy (87%), most had metastatic disease at study entry (91%), and almost half had failed prior anthracycline-based chemotherapy (48%).

**Table 1.** Demographics and clinical characteristics of patients enrolled in the SQ3370-001 Phase 1 dose-escalation trial

| Characteristic | Phase 1 Cohort |
| --- | --- |
|  | N = 23<sup>a</sup> |
| Median age in years (range) | 59 (26–92) |
| Male, n (%) | 11 (48) |
| White, n (%) | 20 (87) |
| ECOG<sup>b</sup> 1, n (%) | 15 (65) |
| Prior surgery, n (%) | 20 (87) |
| Prior radiotherapy, n (%) | 11 (48) |
| Prior systemic anti-cancer therapy, n (%) | 20 (87) |
| Median number of prior systemic anti-cancer therapy (range) | 4 (0–14) |
| Received prior anthracycline, n (%) | 11 (48) |
| Diagnosis of sarcoma,<sup>c</sup> n (%) | 15 (65) |
| Metastases at study entry, n (%) | 21 (91) |

<sup>a</sup>Number of patients who received SQ3370 up to the 12x dose level. Only 22 patients were evaluable for DLT (1 patient in the 12x cohort who developed COVID-19 Day 2/Cycle 1 was excluded). <sup>b</sup>Patients with ECOG 0 and 1 were eligible. <sup>c</sup>Other solid tumors included: breast cancer (*n* = 2); gynecological cancers (*n* = 2); inflammatory myofibroblastic tumor (*n* = 1), Merkel cell carcinoma (*n* = 1), squamous cell carcinoma of the supraclavicular fossa (*n* = 1) and solitary fibrous tumor (*n* = 1). 1x = 75 mg/m<sup>2</sup> of Dox Eq. Dox = doxorubicin; Dox Eq = doxorubicin molar equivalents; ECOG = Eastern Cooperative Oncology Group.

#### SQ3370 shows evidence of safety and tolerability in patients

The median (range) number of SQ3370 treatment cycles received was 3 (1–18). Of note, one patient receiving 6x the conventional Dox dose completed 18 cycles of SQ3370 treatment and was on the study for over a year.

An overview of notable treatment emergent adverse events (TEAEs) in dose cohorts of SQ3370 is shown in Table 2. Comparable to the data seen in preclinical studies, SQ3370 treatment was safe and tolerable in patients across the evaluated dose levels.

**Table 2.**
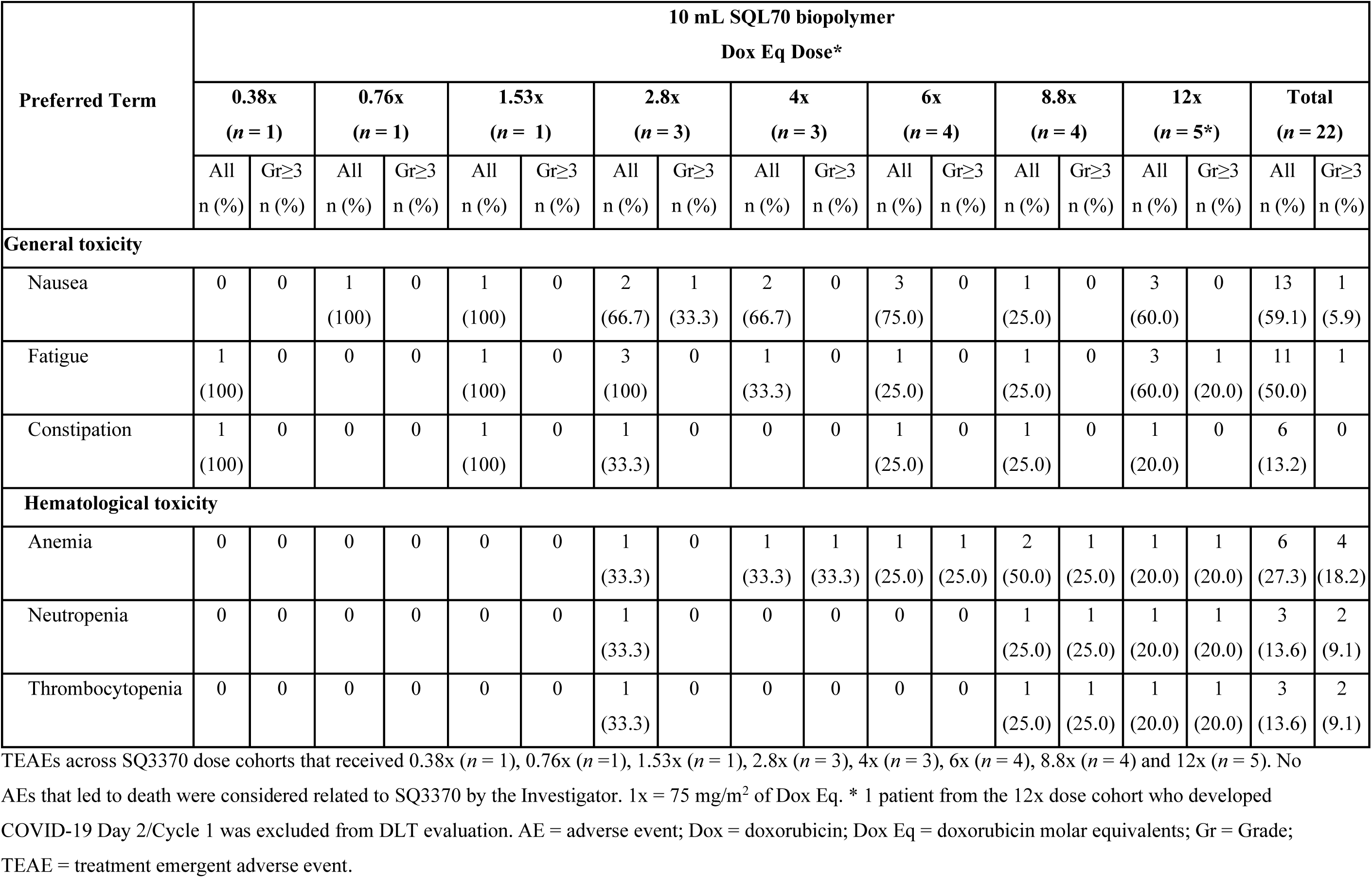
TEAEs across SQ3370 dose cohorts in patients in the SQ3370-001 Phase 1 trial

| Preferred Term | 10 mL SQL70 biopolymer<br>Dox Eq Dose* |  |  |  |  |  |  |  |  |  |  |  |  |  |  |  |  |  |
| --- | --- | --- | --- | --- | --- | --- | --- | --- | --- | --- | --- | --- | --- | --- | --- | --- | --- | --- |
|  | 0.38x<br>(n = 1) |  | 0.76x<br>(n = 1) |  | 1.53x<br>(n = 1) |  | 2.8x<br>(n = 3) |  | 4x<br>(n = 3) |  | 6x<br>(n = 4) |  | 8.8x<br>(n = 4) |  | 12x<br>(n = 5*) |  | Total<br>(n = 22) |  |
|  | All<br>n (%) | Gr≥3<br>n (%) | All<br>n (%) | Gr≥3<br>n (%) | All<br>n (%) | Gr≥3<br>n (%) | All<br>n (%) | Gr≥3<br>n (%) | All<br>n (%) | Gr≥3<br>n (%) | All<br>n (%) | Gr≥3<br>n (%) | All<br>n (%) | Gr≥3<br>n (%) | All<br>n (%) | Gr≥3<br>n (%) | All<br>n (%) | Gr≥3<br>n (%) |
| **General toxicity** |  |  |  |  |  |  |  |  |  |  |  |  |  |  |  |  |  |  |
| Nausea | 0 | 0 | 1<br>(100) | 0 | 1<br>(100) | 0 | 2<br>(66.7) | 1<br>(33.3) | 2<br>(66.7) | 0 | 3<br>(75.0) | 0 | 1<br>(25.0) | 0 | 3<br>(60.0) | 0 | 13<br>(59.1) | 1<br>(5.9) |
| Fatigue | 1<br>(100) | 0 | 0 | 0 | 1<br>(100) | 0 | 3<br>(100) | 0 | 1<br>(33.3) | 0 | 1<br>(25.0) | 0 | 1<br>(25.0) | 0 | 3<br>(60.0) | 1<br>(20.0) | 11<br>(50.0) | 1 |
| Constipation | 1<br>(100) | 0 | 0 | 0 | 1<br>(100) | 0 | 1<br>(33.3) | 0 | 0 | 0 | 1<br>(25.0) | 0 | 1<br>(25.0) | 0 | 1<br>(20.0) | 0 | 6<br>(13.2) | 0 |
| **Hematological toxicity** |  |  |  |  |  |  |  |  |  |  |  |  |  |  |  |  |  |  |
| Anemia | 0 | 0 | 0 | 0 | 0 | 0 | 1<br>(33.3) | 0 | 1<br>(33.3) | 1<br>(33.3) | 1<br>(25.0) | 1<br>(25.0) | 2<br>(50.0) | 1<br>(25.0) | 1<br>(20.0) | 1<br>(20.0) | 6<br>(27.3) | 4<br>(18.2) |
| Neutropenia | 0 | 0 | 0 | 0 | 0 | 0 | 1<br>(33.3) | 0 | 0 | 0 | 0 | 0 | 1<br>(25.0) | 1<br>(25.0) | 1<br>(20.0) | 1<br>(20.0) | 3<br>(13.6) | 2<br>(9.1) |
| Thrombocytopenia | 0 | 0 | 0 | 0 | 0 | 0 | 1<br>(33.3) | 0 | 0 | 0 | 0 | 0 | 1<br>(25.0) | 1<br>(25.0) | 1<br>(20.0) | 1<br>(20.0) | 3<br>(13.6) | 2<br>(9.1) |

TEAEs across SQ3370 dose cohorts that received 0.38x (n = 1), 0.76x (n = 1), 1.53x (n = 1), 2.8x (n = 3), 4x (n = 3), 6x (n = 4), 8.8x (n = 4) and 12x (n = 5). No AEs that led to death were considered related to SQ3370 by the Investigator. 1x = 75 mg/m<sup>2</sup> of Dox Eq. \* 1 patient from the 12x dose cohort who developed COVID-19 Day 2/Cycle 1 was excluded from DLT evaluation. AE = adverse event; Dox = doxorubicin; Dox Eq = doxorubicin molar equivalents; Gr = Grade; TEAE = treatment emergent adverse event.

Regardless of causality, the most frequently reported any grade TEAEs included nausea (13/22 patients, 59.1%) and fatigue (11/22 patients, 50.0%) (Table 2). The most frequent Grade ≥ 3 was anemia (6/22 patients, 27.3%). Neutropenia rates were low, with 3/22 patients (13.6%) experiencing any grade TEAE neutropenia, with only one Grade ≥ 3 TEAE neutropenia each occurring in the 8.8x and 12x dose cohorts. Of note, patients were allowed to enroll into the study with Grade 1 anemia and only two patients utilized supportive measures (e.g., granulocyte colony stimulating factor). A complete determination of cardiotoxic effects from SQ3370 in humans requires a longer duration of follow-up and therefore is beyond the scope of this manuscript [37, 39].

#### Plasma pharmacokinetics in patients treated with SQ3370 resemble preclinical observations

Plasma PK profiles in patients treated with SQ3370 showed similar trends to those observed previously in animals (Fig. 5). As in rats (Fig. 5A) and dogs (Supplementary Figure S3), SQP33 protodrug on day 1 was cleared rapidly from the plasma of patients due to capture by SQL70 biopolymer, with an associated increase in circulating plasma Dox (Fig. 5B). Preliminary analysis over multiple dose levels showed that overall plasma exposure to Dox (AUC over the 5-day dosing period) generally increased with SQ3370 dose level (Supplementary Table S1). Dox AUC levels for doses greater than 2.8x were higher than those clinically observed for conventional Dox [37, 40]. However, the Dox maximum concentration (C_max_) values in the plasma on day 1 of the 5-day regimen remained below those predicted for conventional Dox [37, 40]. Taken together, the data suggest that SQ3370, even at 12x, can expand the systemic exposure (AUC) of Dox beyond what was previously possible, with a C_max_ below that of conventional Dox.

**Fig. 5.**
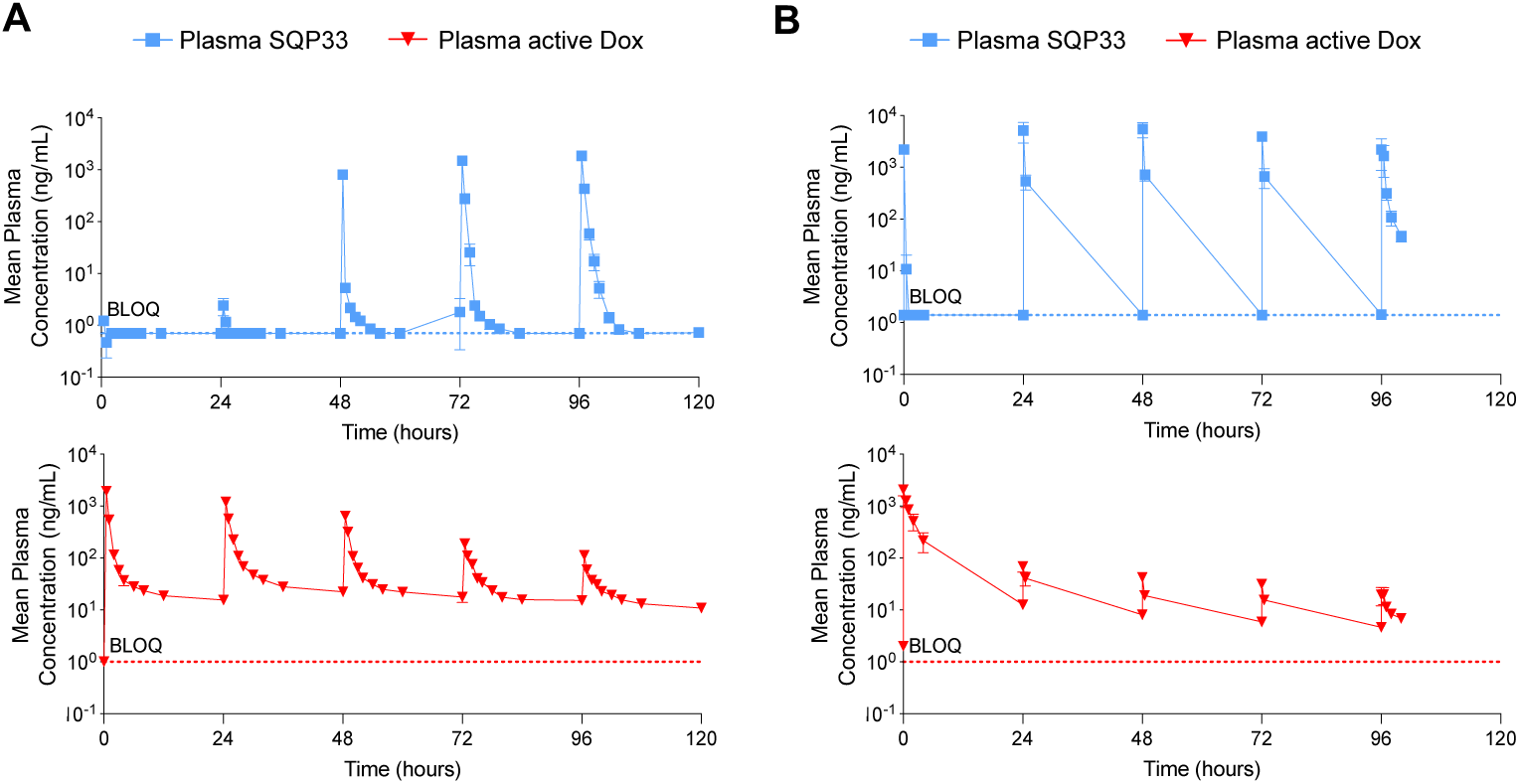
Plasma PK profiles in rats and patients with advanced cancer. (**A**)-(**B**) Five-day plasma concentration-time curves of SQP33 protodrug or Dox in (**A**) rats (*n* = 3, cumulative dose of 107.5 mg/kg/cycle Dox Eq) [29] and (**B**) patients from the SQ3370-001 Phase 1 trial treated with SQ3370 (*n* = 3, 4x dose level = 304 mg/m^2^ Dox Eq, where 1x = 75 mg/m^2^ of conventional Dox). SQL70 biopolymer was given SC (rats) or intratumorally (cancer patients) on day 1, followed by SQP33 protodrug given IV once daily for 5 days. Shown are mean ± SEM. BLOQ = below the limit of quantification; conc = concentration; Dox = doxorubicin; Dox Eq = doxorubicin molar equivalents; IV = intravenous; PK = pharmacokinetics; SC = subcutaneous, SEM = standard error of mean.

#### SQ3370 induces pharmacodynamic T-cell dependent immune activation in the tumor microenvironment in patient tumor biopsies

Immune cell profiling of tumor biopsies from patients who received greater than the 2.8x Dox dose level is presented in Fig. 6. After one cycle of SQ3370, patient tumor biopsies showed an increase in cell density of Granzyme B^+^ CD3^+^ and Granzyme B^+^ CD8^+^ T-cells, suggesting elevated cytotoxic lymphocyte activity in tumors (Fig. 6A-B). This increase in cytotoxic activity correlated with a trend of increase in tumor cell apoptosis confirmed by cleaved caspase-3 (CC3) staining (Supplementary Figure S4). These observed effects on the tumor immune microenvironment build on the previous preclinical findings of increased T-cell infiltration into tumors following SQ3370 treatment in a mouse model [28].

**Fig. 6.**
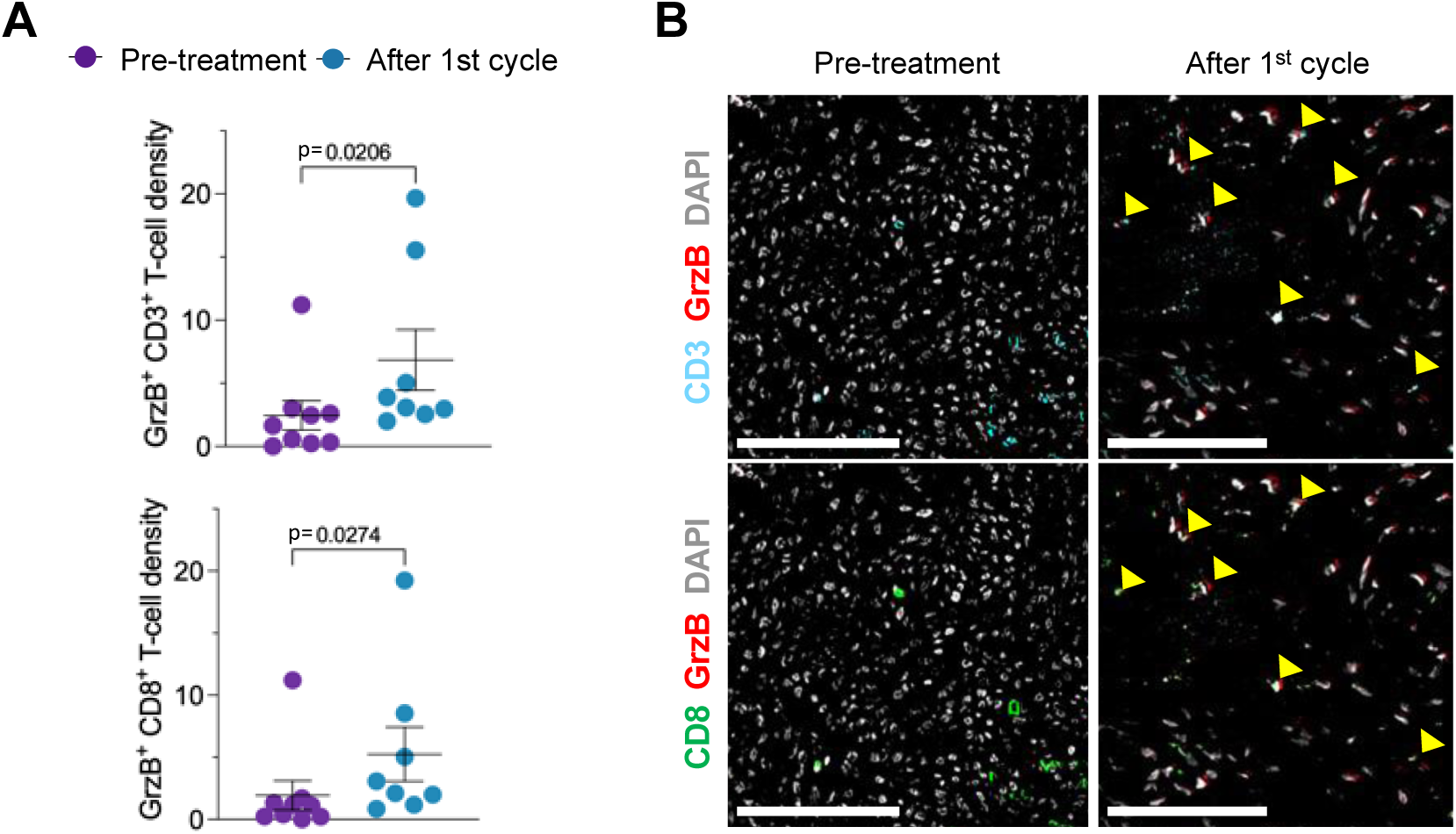
SQ3370 leads to activation of cytotoxic T cells in patient tumor biopsies. (**A**) Tumors from patients in SQ3370-001 clinical trial dose cohorts 4x (*n* = 3), 6x (*n* = 3), and 8.8x (*n* = 4) were analyzed by mIHC pre-treatment (baseline) and at 21 days after initiation of the first cycle of SQ3370 treatment. Increased cytotoxic activity of CD3^+^ and CD8^+^ T-cells was assessed by GrzB expression. Shown are mean ± SEM. P-values were determined by Mann-Whitney U. (**B**) Panel of mIHC images containing two merged pseudo-fluorescent colors of one region from one patient (at the 8.8x dose level) collected pre-treatment and after completion of one SQ3370 treatment cycle in the SQ3370-001 clinical trial. Top panel images are merged pseudo-color stains for CD3 and GrzB. Bottom panel images are for the same region with merged pseudo-colors stains for CD8 and GrzB. Yellow arrows point to double-positive cells. Gray shows all cells by DAPI staining. Scale bars: 100 µm. DAPI = 4′,6-diamidino-2-phenylindole; GrzB = granzyme B; mIHC = multiplexed immunohistochemistry; SEM = standard error of mean.

## Discussion

The CAPAC platform is designed to activate cancer therapies at the tumor, increasing their TI and unlocking new biological effects. We have previously reported on the chemistry of SQ3370 along with its preclinical safety, PK, and efficacy in rodents [28, 29]. Those studies showed reduced Dox cytotoxicity in vitro, and potent anti-tumor effects in animal models. In this article, we demonstrate SQ3370’s ability to localize previously unachievable levels of Dox to mouse tumors. Further, we show translatability of preclinical data (safety, toxicological, and PK observations) through to the clinic as well as SQ3370’s ability to activate the immune system in patient tumors.

By activating Dox directly at the tumor site — confirmed by imaging (Fig. 2) and functional assessments (Supplementary Figure S1) — it was possible to administer unprecedented dose levels of Dox with manageable side effects, regardless of the species. SQ3370 delivered 19.1 times the MTD of conventional Dox in mice and 10.8 times the MTD of conventional Dox in rats [29], with no MTD reached to date. The enhanced tolerability of SQ3370 at high doses in rodent [29] and dog models translated to patients in the clinic, with patients receiving up to 12x the approved clinical dose of Dox per cycle, with no DLTs reported. TEAEs with the high levels of Dox delivered by SQ3370 were minimal, with 3 patients experiencing Grade ≥ 3 anemia and 1 patient experiencing Grade ≥ 3 neutropenia. This is an improvement from the well characterized and severe side effects of conventional Dox in patients, including neutropenia, thrombocytopenia, and anemia [37]. The unprecedented doses of Dox possible with SQ3370 and limited TEAEs observed confirm the exquisite selectivity of protodrugs that can be activated in humans by click chemistry.

In terms of cardiotoxicity of SQ3370, long term follow up is needed. The short period of SQ3370 treatment in patients, up to > 1 year in a patient receiving 6x the conventional Dox dose, is not sufficient to evaluate cardiotoxic effects. In current clinical practice, most clinicians limit the cumulative dose of Dox to 400–450 mg/m^2^ [26–29], but considerable cardiac damage is known to occur at dosages considerably below this level [41, 42]. To date, 19 out of 22 evaluable patients (86.4%) in the Phase 1 dose escalation cohorts have received > 400 mg/m^2^ cumulative Dox given as SQP33 protodrug. While cardiotoxicity did not limit dose escalation during the study, serious, irreversible heart problems can emerge months after cessation of treatment. Continuous evaluation of patients in the Phase 1 study and in subsequent studies is required to understand the long-term effects of SQ3370 on cardiac function.

The main benefit of therapeutics activated at the site of the tumor is that they may eliminate cancerous cells while sparing healthy tissue from cytotoxic effects, which may, in part, account for the low rates of adverse events with SQ3370. Previous approaches with locally activated therapeutics have attempted to use inherent biological differences between cancerous and healthy tissue such as enhanced vascular permeability/poor lymphatic drainage (aldoxorucibin and doxil) [10, 43–45], oxygen level [12], or protein expression [13]. Some therapeutics designed to be cleaved by proteases elevated in cancerous cells have been developed by modifying the aminoglycoside portion of Dox with peptides such as (i) L-377202 activated by the prostate-specific antigen enzyme [46–48], (ii) DTS-201 (CPI-0004Na) activated by endopeptidases [13, 49], and (iii) “Compound 5” activated by matrix metalloproteinases [50]. Generally, these approaches failed to enhance the TI of Dox, while other approaches were limited by poor translatability from preclinical models to humans. The consistently higher Dox doses administered across species with SQ3370 go far beyond what has been previously possible with Dox using either conventional or biology-dependent targeted approaches [13, 43–47, 49–51] (Supplementary Figure S5). Relying on chemistry rather than biological factors has allowed SQ3370 to overcome the limited translatability of preclinical findings to humans.

The tolerability of SQ3370 is also consistent with the PK characteristics observed across species. Previous studies in rodents [29] and data reported here from dogs and patients indicate that locally injected SQL70 biopolymer effectively captures circulating SQP33 protodrug, followed by rapid activation and release of Dox over the course of the 5-day dosing regimen. The plasma concentration-time curves in humans showed consistency with PK data from animals (Fig. 5). The key difference between the animal and human PK studies was that the animals were non-tumor bearing. Assessment of biopolymer distribution and its impact on PK is currently being evaluated in tumor-bearing animal models.

In patients treated with SQ3370, the AUC of active Dox exceeded the AUC of conventional Dox [37, 40] at ≥ 4x dose level. Conversely, even at the highest dose level (12x) of SQ3370, the C_max_ of Dox did not exceed that of conventional Dox [37, 40]. Despite the high AUCs measured in patients, SQ3370 was well tolerated at dose levels presented here including the highest dose (12x). While this is consistent with some literature reports on conventional Dox suggesting that acute effects including nausea, vomiting, and hematological side effects are lessened by a decrease in C_max_ [52–54], it contrasts with other studies that have shown significant increases in some toxicities such as mucositis resulting from higher AUCs [55, 56]. Evidently, the relationship of PK and toxicity of conventional Dox is not fully understood in the literature. As the SQ3370 clinical study proceeds, the interplay of Dox PK and toxicity will be evaluated further with long term patient follow-up.

One of the limitations of both the preclinical and human PK data is that active Dox levels were measured systemically in the plasma, and not directly at the site of the tumor due to technical limitations associated with capturing tumor samples for analysis. This approach does not distinguish Dox that was released locally at the site of the tumor and entered systemic circulation, from Dox released into the circulation directly.

The presence of active systemic Dox is likely a combination of locally released Dox that diffuses from the tumor into systemic circulation and biopolymer that escapes from the tumor which in turn activates the protodrug encountered in circulation. Regardless of the mechanism, systemic exposure (AUC) of Dox beyond what was previously possible with conventional Dox suggests clinical benefit of SQ3370 in being able to treat not only local disease, but also micrometastatic or widely disseminated lesions that may not be accessible to intratumoral injections. We previously reported SQ3370’s ability to induce sustained anti-tumor responses against biopolymer-injected and non-injected distal tumors in an MC38, MCA-205, B16-F10 syngeneic tumor models [28, 57]. While SQ3370’s tumor growth inhibition was significantly better in the locally-injected MC38 tumor shown here, the effect on the distal lesion was at least as good as that of conventional Dox (Supplemental Figure S1). This anti-tumor effect on the distal lesion was likely due to systemically circulating Dox following SQ3370 treatment.

While Dox is capable of immune activation via immunogenic cell death in restricted *in vitro* environments [58, 59], its narrow TI and the associated systemic immunosuppression make this effect challenging to translate *in vivo*. The promise of CAPAC is that by safely administering unprecedented doses of Dox directly at the tumor site, thereby limiting immunosuppression, some of the immune activation effects can be unlocked *in vivo*. Imaging with MALDI provided a snapshot of the high concentration and spatial distribution of Dox exposure within the tumor and stroma after SQ3370 treatment and the subsequent advanced necrosis seen in the tumor. In essence, we obtained visual proof that SQ3370 enables levels of Dox exposure at the tumor that would have been unimaginable prior to use of the CAPAC platform.

Associated with the high levels of Dox in patients, we observed anti-tumor immune activation with a shift towards a T-cell supportive tumor immune microenvironment and increased numbers of functional cytotoxic T cells (granzyme B^+^ CD3^+^ or CD8^+^). It is possible that in the absence of the characteristic myelosuppression associated with systemic administration of conventional Dox, the high doses of Dox locally delivered by SQ3370 treatment can result in activation of the immune system. Data from patient tumor biopsies reported here showed increased tumor necrosis within the tumor and tumor microenvironment after SQ3370 treatment. Taken together, these observations suggest that the localization of high doses of Dox using SQ3370 may activate the immune system and lead to tumor cell death. This ability of SQ3370 to activate the immune system could also benefit patients with micrometastatic disease.

The immune activation in patient tumor biopsies was even more impressive because most patients in the trial had refractory disease, were heavily pre-treated, and, as a result, likely had exhausted immune systems prior to SQ3370 treatment. Furthermore, even patients with soft tissue sarcomas (*n* = 11, 65%), a cancer type that is poorly responsive to checkpoint inhibition therapies and characterized by microenvironments that are poorly immunogenic [60], experienced immune activation. SQ3370 could be beneficial as monotherapy and ideally suited to use in combination with checkpoint inhibitors or other immune modulators.

The CAPAC platform has the potential to transform cancer treatment through several specific benefits. First, the use of click chemistry eliminates the dependency on biological factors for conditional activation; but rather, the activation of the protodrug is based on a chemical reaction that is fast, specific, and efficient. As discussed in this article, this also improves translatability across species. Second, by using click chemistry to activate the drug at the site of the tumor, high doses of toxic cancer drugs can be administered as attenuated protodrugs. This approach may expand the use of drugs that were not viable systemic therapies on their own due to their toxicity (e.g., monomethyl auristatin E). Finally, decoupling the targeting agent from the protodrug allows the creation of a modular system with the potential to enable cancer drug combinations, activated at the tumor with off-the-shelf payloads, previously prevented by overlapping toxicities.

We are currently building a library of protodrug therapeutics based on commonly used chemotherapeutic agents. We are simultaneously developing the next generation of CAPAC with tumor-targeting agents that can be administered systemically and become localized at the site of the tumor via antigen targeting. These therapies can provide a more effective alternative to antibody-drug conjugates that can often be limited by narrow TI, off-target toxicity, and undesirable immunogenic effects from aggregation [14, 16]. Studies exploring a HER2-targeted approach and an MMAE payload are ongoing (manuscript in preparation).

## Conclusion

SQ3370, the first example of a drug developed within the CAPAC platform, showed consistent PK, safety, and toxicology profiles across preclinical studies, which translated to similar findings in a Phase 1 first-in-human clinical study despite a heavily pretreated, relapsed, and refractory patient population. By activating unprecedented levels of Dox directly at the tumor, SQ3370 has the potential to expand the TI of Dox and unlock new biological effects, such as novel immune activation. These aspects are being further explored in the ongoing Phase 2a part of the trial, which will further evaluate safety, tolerability, and the ability of SQ3370 to improve response rates in Dox-naïve patients.

## Abbreviations

AUC: Area Under the Curve
CAPAC: Click Activated Protodrugs Against Cancer
CC3: Cleaved caspase-3
C_max_: Maximum Concentration
DLT: Dose-limiting Toxicity
Dox: Doxorubicin
Dox Eq: Doxorubicin molar equivalents
H&E: Hematoxylin and Eosin
HNSTD: Highest Non-Severely Toxic Dose
MALDI-IMS: Matrix-assisted Laser Desorption/ionization-Imaging Mass Spectrometry
MTD: Maximum Tolerated Dose
NOAEL: No Observed Adverse Effect Level
RP2D: Recommended Phase 2 Dose
TEAE: Treatment Emergent Adverse Event
TI: Therapeutic Index
TILs: Tumor Infiltrating Lymphocytes
TK: Toxicokinetics
TME: Tumor Microenvironment
WBC: White Blood Count

## Declarations

### Ethics approval and consent to participate

Animal studies were conducted under the protocols and/or guidelines approved by the Institutional Animal Care and Use Committee (IACUC) of Explora BioLabs (San Diego, CA, USA), Cephrim Biosciences, Inc. (Woburn, MA, USA) or Covance Laboratories (Madison, WI, USA). The SQ3370-001 Phase 1 trial was conducted under the clinical study protocol approved by the central WCG Institutional Review Board (IRB), local IRBs or Ethics Committees (ECs) at the listed clinical sites: Stanford Cancer Center, MD Anderson Cancer Center, Henry Ford Hospital, Mary Crowley Cancer Research, Washington University St. Louis, Sarcoma Oncology, Royal North Shore Hospital, Chris O’Brien Lifehouse and Cancer South Australia.

### Consent for publication

Not applicable

### Availability of data and materials

The data generated in this study are available within the article and its supplementary data files.

### Authors’ contributions

Study conceptualization: SS, NAY, MA, T-HN, and JMMO. Data curation: SS, NAY, MZ, MA, and T-HN. Formal analysis: SS, NAY, MA, AM, and T-HN. Funding acquisition: SS and JMMO. Investigation: SS, NAY, MA, AM, and JMMO. Visualization: SS, NAY, MZ, MA, AM, and JMMO. Writing manuscript drafts: SS, NAY, MZ, MA, T-HN, and JMMO. All authors reviewed drafts and have read and approved the final draft of the manuscript.

## Acknowledgments

The authors thank all patients and their families for participating in SQ3370-001. The authors also thank the clinical and preclinical research teams for generating the data (Bryan Calaway, Jim Williams, M.D., and Matthew Tso,) and for critical review of the manuscript (Steve Abella, M.D., Scott Wieland, Ph.D., Sadie Whittaker, Ph.D., and Jesse McFarland, Ph.D.). Medical writing support was provided by Martha Mutomba, on behalf of Shasqi Inc.

