## Supplemental Information for "SQ3370, the first clinical click chemistry-activated cancer therapeutic, shows safety in humans and translatability across species"

**Title**

**Authors and Institutions**

Sangeetha Srinivasan[1], Nathan A. Yee[1], Michael Zakharian[1], Maša Alečković[1], Amir Mahmoodi[1], Tri-Hung Nguyen[1], José M. Mejía Oneto[1]

[1]Shasqi Inc, 665 3rd St, Suite 501, San Francisco, CA 94107, USA

**Corresponding Author**

José M. Mejía Oneto, MD, PhD; Shasqi Inc., 665 3rd St, Suite 501, San Francisco, CA 94107, USA.

**Target Journal:** ***BMC Molecular Cancer***

**Supplementary Material Table of Contents**

Supplementary Materials and Methods

Preclinical studies

Clinical trial data

Supplementary Results

Preclinical studies

Supplementary Figure S1. Peritumoral injection is superior to distal injection of SQL70

Supplementary Figure S2. Tumor Dox activation and distribution after control treatments in a mouse tumor model

Supplementary Figure S3. Pharmacokinetics of Dox activation with SQ3370 in dogs

Supplementary Figure S4. SQ3370-001 patient tumor biopsies show trends of increased tumor cell apoptosis after 1 cycle of SQ3370 treatment

Supplementary Figure S5. Click chemistry surpasses translatability of previous Dox-based approaches

Supplementary Table S1. Cycle 1 SQ3370 – Active Dox Cmax at 5 minutes on day 1 and 5-day AUC values in patients

### Supplementary Materials and Methods

#### *Preclinical studies*

*Plasma sample preparation for TK assessment in dogs*

Sample preparation for LC-MS/MS: To quantify SQP33 in dog plasma, a solid-phase extraction method was used. In brief, 50.0 µL of sample plasma was transferred to a low-binding 96-well plate. Standard curve calibrators, QCs, blank plasma, and reagent blank were added to the corresponding wells on the 96-well plate. 50 µL of internal standard (IS) containing 1,000 ng/mL SQP33 structural analog in water:acetonitrile (50:50) was added to all samples except matrix blanks and mixed well. 100 µL of 100 mM ammonium acetate was then added to all samples, mixed, and centrifuged at 3,000 rpm for 1 minute. After the centrifugation, 200 µL of each sample was transferred to the SPE plate (Oasis HLB µ Elution Plate, Waters), which was previously prepared according to the product manual. The SPE plate was washed once with 200 µL of 100 mM ammonium acetate. Finally, the SPE plate is eluted with 25.0 µL 5% formic acid (FA)-methanol into a low-binding plate containing 150 µL 1% CHAPS in water for LC-MS/MS analysis. To quantify Dox in dog plasma, a protein precipitation method was used. 50.0 µL of sample plasma was transferred to a low-binding 96-well plate (including calibrators, Q, Cs, unknown samples, blank dog plasma, and reagent blank). 50.0 µL of IS working solution [200 ng/mL Dox-$^{13}$C-$d_3$ in water:acetonitrile (50:50)] was added to all samples except matrix blanks, followed by centrifugation at 3,000 rpm for 1 minute. 20 µL of 10% FA in water was added, mixed, followed by 500 µL of acetonitrile to all samples, and vortexed for 5 min. The samples were centrifuged at 4,000 rpm for 5 min, then transferred 450 µL of the supernatant to a new 96-well plate. Samples were dried down at 40 °C under nitrogen in the Turbovap. The samples were reconstituted in 200 µL of 90:10 water: acetonitrile for LC-MS/MS analysis.

*LC-MS/MS analysis*

Analytical HPLC was performed using an Xbridge Phenyl column (50 x 2.0 mm, 5.0 µm particle size, Waters Corporation, Milford, MA). An injection volume of 20.0 µL for SQP33, or 10.0 µL for Dox, was used with a flow rate of 0.4 mL/min, and a gradient mobile phase of 0.1% FA-water and 0.1% FA-acetonitrile. Mass spectrometry (SCIEX QTRAP 6500 (SCIEX, Foster City, CA) for SQP33 analysis; SCIEX API 5000 for Dox analysis) was connected in tandem with the HPLC. ESI+ ionization was used for Dox analysis with the cone voltage of 5,100V and 500 ˚C, while SQP33 was analyzed in ESI- ionization mode with the cone voltage and temperature of 4,500V and 200 ˚C respectively. Transitions monitored were as follows: Dox: 544.1 –> 361.3 m/z; Dox-$^{13}C$-$d_3$: 548.2 –> 401.1 m/z; SQP33: 809.2 –> 413.2 m/z; SQP33-IS-Analog: 823.4 –> 427.7 m/z. The SQP33 IS analog used for these studies was TCO-Dox-Alanine (Scheme 1), a structural analog of SQP33.

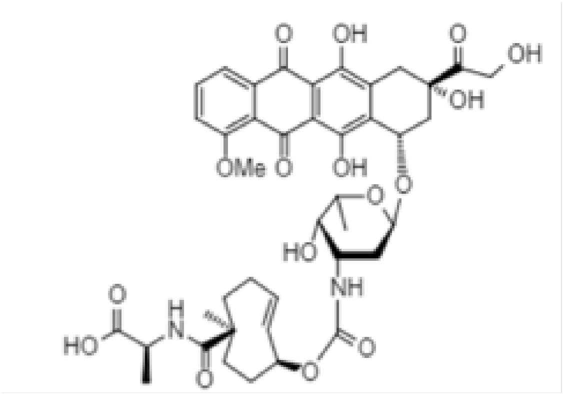


**Scheme 1:** TCO-Dox-Alanine

*Tissue slides preparation for MALDI MS-Imaging and hematoxylin and eosin (H&E) staining*

Mouse tumor tissues were harvested according to the approved IACUC protocol. The tumor was cryo-sectioned at a chamber temperature of -20 ºC with a 10 µm thickness setting. Sections were thaw-mounted onto the ITO-coated side or Superfrost slides for mass spectrometry imaging (MSI) and H&E staining, respectively. Slides were stored in the -80 ºC freezer until the time for imaging. To prepare the slide for imaging, tissue sections on ITO slides were transferred from -80 ºC to a vacuum desiccator and dried for 15 minutes before matrix deposition. An optical image of tissue sections was obtained using a scanner to synchronize the positions of the tissue sections with the target of the laser. Once the optical image was obtained, the slide was immediately transferred to the HTX TM-Sprayer (HTX Technologies, Chapel Hill, NC) for matrix deposition. DAN (1,5-diaminonaphtalene) was applied on the tissue sections at 10 mg/mL in 1:1 acetonitrile:water.

MALDI-MS imaging was performed in continuous accumulation of selected ions (CASI) mode and negative ionization over a mass range of m/z 397 +/-30 Da for Dox and SQP33 protodrug (in-source fragments) on a Bruker Solarix (MALDI-FTICR) mass spectrometer. Imaging was performed using a laser raster size of 70 µm custom setting for the entire tumor section. Data were acquired and analyzed with FlexImaging v5.0 (Bruker). MSI data were analyzed using MultImaging™ v1.2. imaging software package (ImaBiotech, Billerica, MA).

H&E were used to stain adjacent sections to overlay the molecular images with the histological structures. H&E staining was imaged with ImageScope v1.12, and annotations were performed by an experienced histologist (not a certified pathologist).

#### ***Clinical trial data***

*Pharmacokinetic assessment of clinical samples, sample preparation and LC-MS/MS*

Blood was collected from subjects on each day of dosing to characterize the plasma pharmacokinetics of SQP33 protodrug and Dox. Approximately 6.0 mL blood was collected for each timepoint into $K_2$EDTA tubes from which plasma was isolated. On Days 1 and 5, blood was drawn pre-infusion, and at 5 min, 30 min, 1 h, 2 h, and 4 h post-infusion. On Days 2-4, blood was drawn pre-infusion, and at 5 min and 30 min post-infusion. Plasma samples were processed using solid-phase extraction followed by analysis with high performance liquid chromatography-tandem mass spectrometry (LC-MS/MS) to determine SQP33 and Dox concentrations.

Human plasma samples were processed using similar extraction protocols as those used for the dog samples, using a solid-phase extraction method. 50 µL of a solution of *trans*-cyclooctene-amine HCl (0.05 mg/mL in water: acetonitrile, 75:25), as a tetrazine quencher, was added to all samples pre-extraction. Following processing, the samples were analyzed by LC-MS/MS. Analytical HPLC was performed using a Kinetex biphenyl column (100 Å 2.6 µm, 4.0 x 2.0 mm, Phenomenex, Torrance, CA) to separate the analytes. The eluates were monitored by an API4000 MS/MS (SCIEX, Foster City, CA) detector in negative MRM mode for SQP33 and positive ion mode for Dox. Dox IS: Dox-$_{13}$C-d$_3$; SQP33 IS: SQP33-$^{13}C_2$-$^{15}$N-d$_2$. The data were acquired and integrated by the data acquisition system Analyst® (SCIEX, Foster City, CA) linked directly to the API4000 MS/MS detector and then processed in Watson LIMS™ (Thermo Scientific).

### Supplementary Results

#### *Preclinical studies*

*GLP toxicology*

All animals survived to their scheduled sacrifice. No SQP33-related, SQL70-related, or SQP33 + SQL70-related effects on clinical, ophthalmic, or macroscopic observations; alterations in body weight, body weight gain, food consumption, or organ weight parameters, or ECG parameters were noted.

The most prominent SQP33 + SQL70-related clinical pathology effects were consistent with decreased hematopoiesis and included moderately or markedly decreased absolute reticulocyte count on day 6 of the dosing phase in males administered SQ3370 at ≥ 3.2 mg/kg/cycle Dox Eq and females administered ≥ 1.8 mg/kg/cycle Dox Eq and minimally decreased white blood cell and absolute lymphocyte counts on day 6 of the dosing phase in males administered SQ3370 at 8.9 mg/kg/cycle Dox Eq and females administered ≥ 3.2 mg/kg/cycle Dox Eq. These effects correlated with the microscopic finding of sternum and femur bone marrow hypocellularity. The white blood cell and lymphocyte decreases were consistent with the Dox active moiety of the protodrug (SQP33). In addition, minimally decreased inorganic phosphorus concentration was observed on day 6 of the dosing phase in females administered SQ3370 at 8.9 mg/kg/cycle Dox Eq. All SQ3370-related clinical pathology effects exhibited evidence of reversibility by day 13 of the recovery phase. No coagulation effects were observed. SQ3370-related microscopic findings at the end of the dosing phase consisted of minimal to moderate bone marrow (sternum and femur) hypocellularity in animals administered 8.9 mg/kg/cycle Dox Eq and minimal bone marrow (sternum and femur) hypocellularity in one female administered 3.2 mg/kg/cycle Dox Eq. Bone marrow was unremarkable in animals administered 8.9 mg/kg/cycle Dox Eq SQP33 without SQL70 biomaterial. SQL70-related subcutaneous injection site reactions were not related to the dose volume. At the end of the recovery phase, SQ3370-related microscopic bone marrow hypocellularity had reversed. SQ3370 at 3.2 mg/kg/cycle Dox Eq was established as the no observed adverse effect level (NOAEL) and 8.9 mg/kg/cycle Dox Eq was established as the HNSTD.

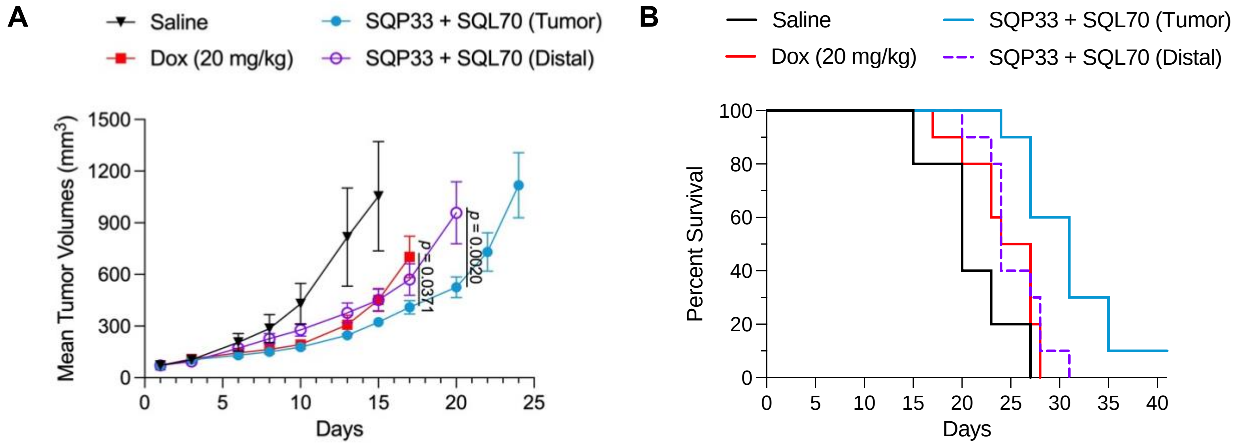


**C**

| Treatments | Median Survival (days) | *p*-value (vs Dox) | Hazard Ratio (logrank); 95% CI of ratio (vs Dox) |
|---|---|---|---|
| Saline | 20 | 0.1137 | 2.041; 0.5715 to 7.289 |
| Dox (20 mg/kg) | 25.5 | N/A | N/A |
| SQP33 + SQL70 (Tumor) | 31 | 0.0031 | 0.3441; 0.1259 to 0.9403 |
| SQP33 + SQL70 (Distal) | 24 | 0.5029 | 0.7896; 0.3266 to 1.909 |

| Treatment | *p*-value (vs SQP33 + SQL70 (Tumor)) | Hazard Ratio (logrank); 95% CI of ratio (vs SQP33 + SQL70 Tumor) |
|---|---|---|
| SQP33 + SQL70 (Distal) | 0.0015 | 0.3789; 0.1825 to 0.7867 |

**Supplementary Fig. S1** Peritumoral injection is superior to distal injection of SQL70. **(A)-(C)** MC38-bearing mice were treated with saline control, Dox (QDx1, IV, 20 mg/kg) or SQL70, implanted peritumorally (Tumor) or at a distant SC location (Distal), followed by SQP33 protodrug infusion, given at 383 mg/kg/cycle Dox Eq (77 mg/kg/day x 5 IV doses from days 1-5). The mean ± SEM of tumor volumes **(A)** and animal survival **(B, C)** are shown ($n$ = 5 mice for saline, $n$ = 10 mice for each of the other groups). *P*-values are relative to the Dox control or SQP33 + SQL70 (Tumor) groups, and only *p*-values ≤ 0.05 are shown in (**A**). *P*-values were calculated by mixed effects analysis **(A)**; Mantel-Cox (logrank) test and logrank Hazard ratios with 95% CIs (**C**). CI = confidence interval; Dox = doxorubicin; Dox Eq = doxorubicin molar equivalents; IV = intravenous; N/A = not applicable; SC = subcutaneous.

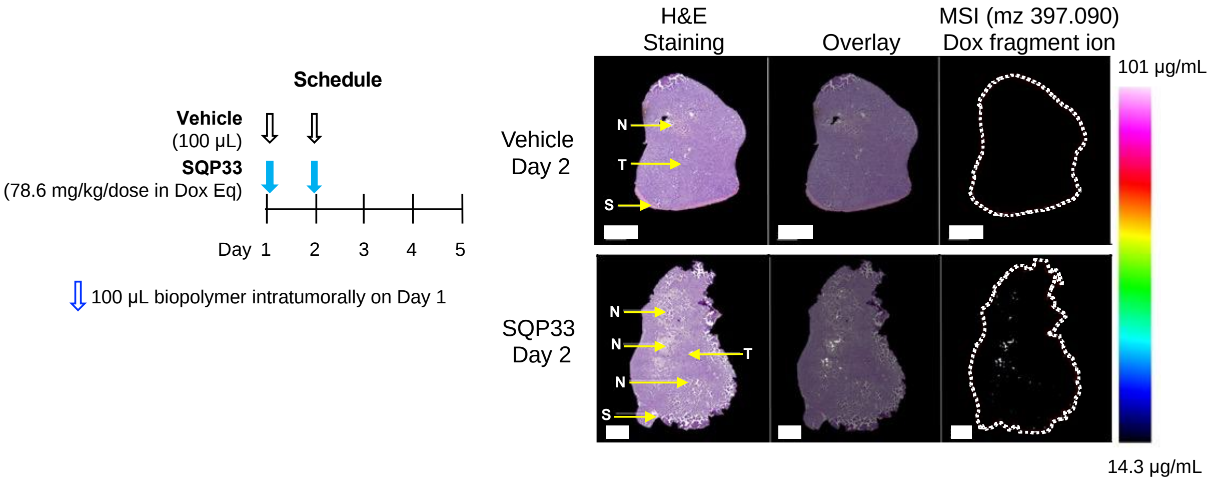


**Supplementary Fig. S2** Tumor Dox activation and distribution after control treatments in a mouse tumor model. Tumor tissues were collected from mice engrafted with MC38 tumor 1 hour after dosing with vehicle or SQP33 (IV, QDx2, 78.6 mg/kg/dose Dox Eq). Serial sections were generated on cryostat for MALDI-MSI and H&E staining. Scale bars: 2 mm. Limit of detection = 9.1 µg/mg of tissue. Dox Eq = Dox molar equivalent; H&E = hematoxylin and eosin; IV = intravenous; MALDI-MSI = matrix assisted laser desorption/ionization mass spectrometry imaging; MSI = mass spectrometry imaging; N = necrosis; S = stroma; T = tumor.

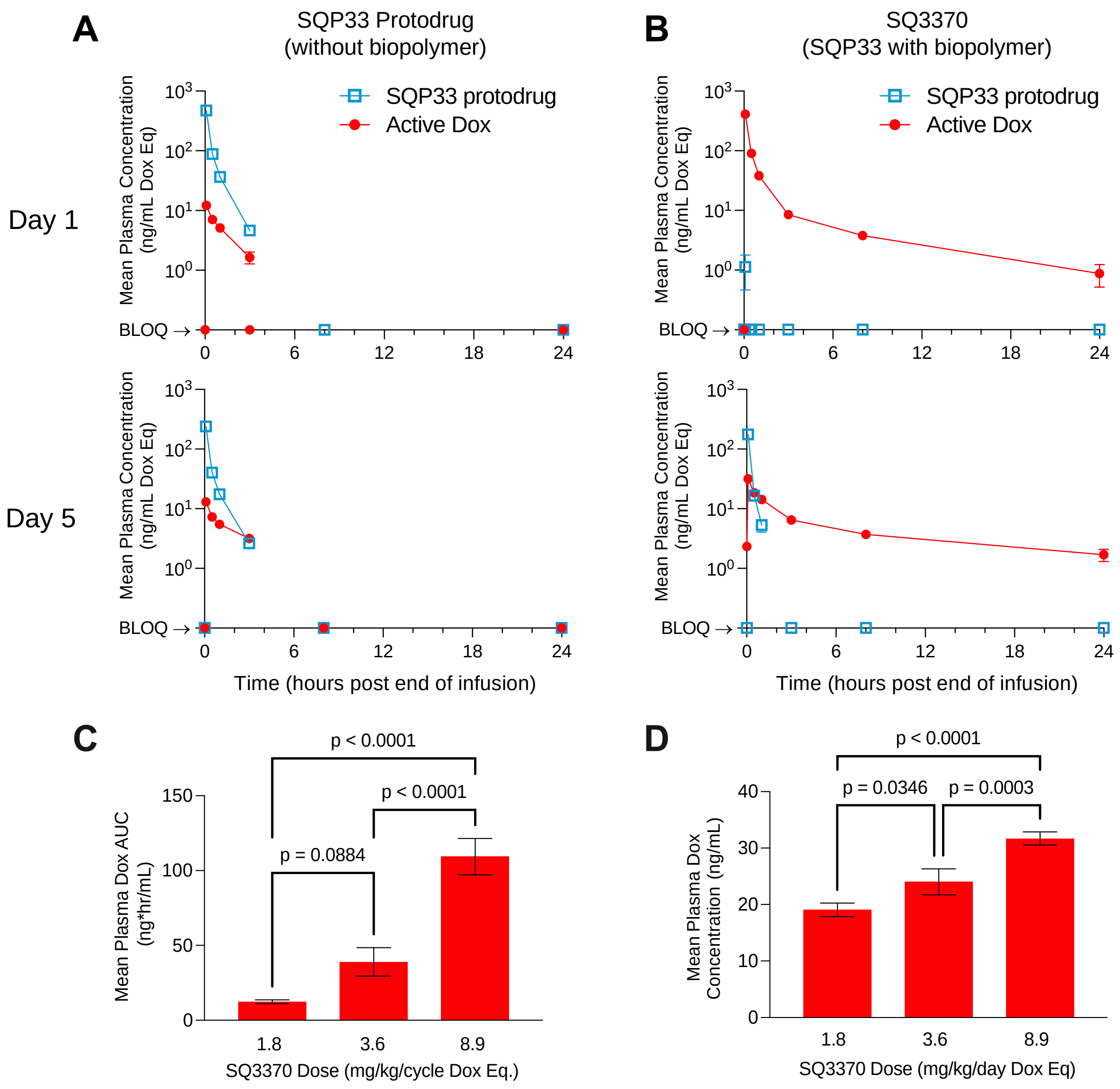


**Supplementary Fig. S3** Pharmacokinetics of Dox activation with SQ3370 in dogs. **(A) – (D)** Plasma concentrations of Dox and SQP33 protodrug on days 1 and 5 after treatment with 5 daily IV doses of SQP33 protodrug at the HNSTD (8.9 mg/kg/cycle Dox Eq) following a 10-mL SC dose of (**A**) vehicle (*n* = 10) or (**B**) SQL70 biopolymer (*n* = 10). SQ3370 doses at 1.8 (*n* = 10), 3.2 (*n* = 6), or 8.9 mg/kg/cycle Dox Eq (*n* = 10) results in dose-dependent (**C**) 24-h AUC and (**D**) $C_{max}$ of plasma Dox concentrations on day 5. Data are shown as mean ± SEM. All groups included a 1:1 ratio of male and female dogs. Statistical analysis done using one-way ANOVA with Tukey's post-test. ANOVA = analysis of variance;

AUC = area under the concentration-time curve; BLOQ = below the limit of quantification (2.00 ng/mL); $C_{max}$ = maximum concentration; dox = doxorubicin; Dox Eq = doxorubicin molar equivalents; HNSTD = highest non-severely toxic dose; IV = intravenous; SC = subcutaneous.

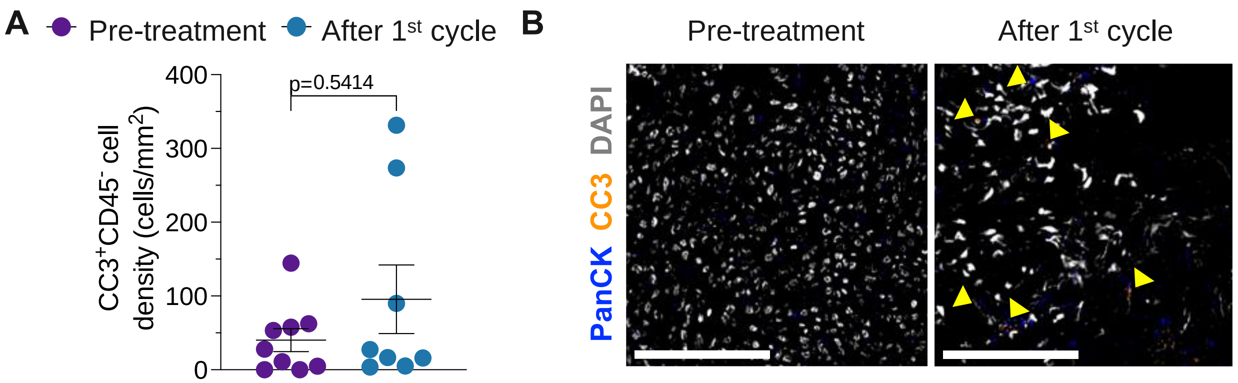


**Supplementary Fig. S4** SQ3370-001 patient tumor biopsies show trends of increased tumor cell apoptosis after 1 cycle of SQ3370 treatment. (**A**) Tumors from patients in SQ3370-001 clinical trial dose cohorts 4x (*n* = 3), 6x (*n* = 3), 8.8x (*n* = 4) were analyzed by mIHC at pre-treatment (baseline) and 21 days after the first cycle of SQ3370 treatment. A trend towards increased apoptosis was observed, marked by $CC3^{+}CD45^{-}$ expression. PanCK expression was not consistent among the patient population (many sarcomas lacked expression) and could therefore not be used for the quantification. *P*-values: Mann-Whitney U. (**B**) Panel of mIHC images containing two merged pseudo-fluorescent colors from one patient (at the 8.8x dose level) collected at baseline and after completion of one SQ3370 treatment cycle in the SQ3370-001 clinical trial. Images are merged pseudo-colors stains for tumor (PanCK) and CC3. Yellow arrows denote double-positive cells. Gray shows all cells by DAPI staining. Scale bars: 100 µm.

CC3 = cleaved caspase 3; DAPI = 4′,6-diamidino-2-phenylindole; mIHC = multiplexed immunohistochemistry; PanCK = Pan cytokeratin.

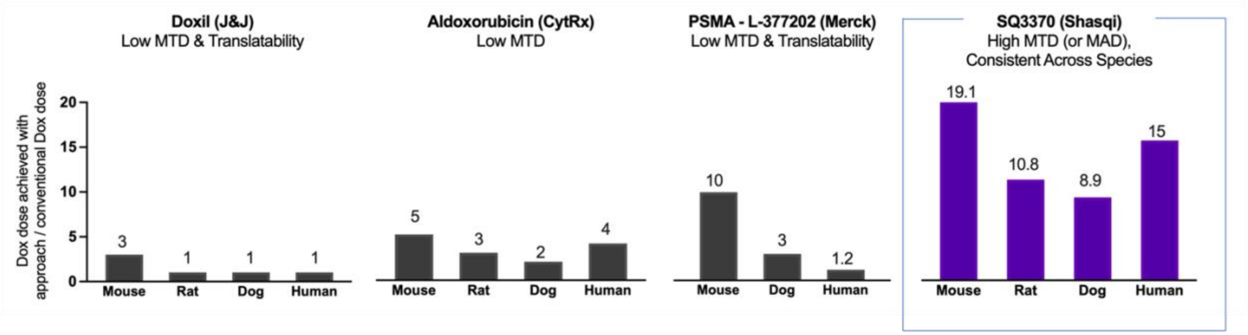


**Supplementary Fig. S5** Click chemistry surpasses translatability of previous Dox-based approaches. The tolerability for several Dox-based products is shown as a ratio of the Dox dose achieved with each approach to the conventional Dox dose in mouse, rat, dog, or human. These include liposomal-based doxil (J&J; Working et al, Hum Exp Toxicol 1996;15(9):751-85 [51]); albumin-binding aldoxorubicin (CytRx; Kratx et al, Hum Exp Toxicol 2007;26(1):19-35 and Uger et al, Clin Cancer Res 2007;13(16):4858-66 [43, 44]); the prodrug requiring enzyme-based activation L-377202 (Merck; DeFeo-Jones et al, Nat Med 2000;6(11):1248-52, DiPaola et al, J Clin Oncol 2002;20(7):1874-9; and Ravel et al Clin Cancer Res 2008;14(4):1258-65 [46-48]); and the click-activated protodrug approach, SQ3370 (Shasqi Inc.). Tolerability is defined by the DLT in each species. For SQ3370: in mice the MTD is 19.1x the single-dose MTD (20 mg/kg) of Dox HCl and in rats, the MTD is 10.8x the multi-dose MTD (2.5 mg/kg) of Dox HCl (Wu et al Chem Sci 2021;12(4):1259-71 [29]); in dogs, the HNSTD of SQ3370 is 8.9x the standard veterinary dose of 1 mg/kg Dox. In humans, SQ3370 has been dosed up to 15x (MAD) the clinical dose of Dox (75 $mg/m^2$). However, the clinical data presented in this article includes up to the 12x dose level only. No MTD has been reached do date. Dox = doxorubicin; J&J = Johnson and Johnson; DLT = dose-limiting toxicity; HNSTD = highest non-severely toxic dose; MAD = maximum administered dose; MTD = maximum tolerated dose.

**Supplementary Table S1** Cycle 1 SQ3370 – Active Dox $C_{max}$ at 5 minutes on day 1 and 5-day AUC values in patients

| **Dox Eq Dose** | **0.38x (*n* = 1)** | **0.76x (*n* = 1)** | **1.53x (*n* = 1)** | **2.8x (*n* = 3)** | **4x (*n* = 3)** | **6x (*n* = 4)** | **8.8x (*n* = 4)** | **12x (*n* = 5[a])** | ***1x Dox Values[b]* (*n* = 23)** |
|---|---|---|---|---|---|---|---|---|---|
| **$C_{max}$** (ng/mL) | 256 | 779 | 203 | 716 ± 547 | 2083 ± 881 | 236 ±222 | 936 ±720 | 865 ±630 | 2570 ±1208 |
| **AUC** (0-$t_{last}$) (ng*h/mL) | 964 | 1621 | 1040 | 2286 ± 1093 | 6677 ±302 | 3733 ±1496 | 6674 ±2212 | 7141 ±3399 | 2580 ±560 **AUC ($0_{h-\infty}$)** |

Cycle 1 SQ3370 $C_{max}$ (5 min. day 1) and 5-day AUC (0-$T_{last}$) for evaluable patients in cohorts with more than one patient (*n* = 11). Patient cohort values show mean ± SD. A single datapoint was extrapolated for a patient in the 8.8x cohort who did not receive the pre-infusion blood draw on day 2. Values for IV Dox are shown as geometric mean ± SD, and AUC shown is from zero to infinity. [a] 1 patient from the 12x dose cohort who developed COVID-19 Day 2/Cycle 1 was excluded from PK evaluation. [b](Source for 1x Dox values in table column in grey color: A. Villalobos et al, Cancer Med 2020;9:882-93 [61] (2060 - 2570 ng/mL); B. Adriamycin product label (2020, Pfizer Inc) [37] (1550-3860 ng*h/mL)). SQ3370-001 data cut off 2022-06-10. AUC = area under the curve; $C_{max}$ = maximum concentration; Dox = doxorubicin; SD = standard deviation; IV = intravenous.
